# Structural mechanism of metal-induced DNA release by an ArsR/SmtB transcriptional regulator

**DOI:** 10.64898/2026.09.30.755606

**Authors:** Diana Meireles, Johnny Lisboa, Rita Pombinho, Ângela Alves, Sandra Sousa, João H. Morais-Cabral, Didier Cabanes

## Abstract

Metal-responsive transcriptional regulators must couple metal coordination to DNA release at the DNA-binding interface. However, the pathway connecting these states remains poorly understood. Here, we define this mechanism for the ArsR/SmtB-family repressor CadC from *Listeria monocytogenes* by combining crystal structures of apo, DNA-bound, and cadmium-bound states with biochemical and *in vivo* analyses. The DNA-bound structure reveals how CadC dimers bind sequence-specifically to operator DNA, via coordinated major-groove, wing and backbone interactions, with high-affinity binding depending on operator architecture. Cadmium is coordinated at the α3N sensor by an unusual Cys3-Met ligand set and stabilizes a DNA-incompatible conformation through remodeling of DNA-contacting elements and steric incompatibility between the ordered N-terminus and DNA. Mutational analyses show how these structural transitions link to metal-dependent transcriptional control in *Listeria* cells. We show that CadC binds zinc and supports zinc-responsive transcription in bacterial cells, and that host zinc availability modifies CadC-dependent infection phenotypes. Together, results reveal how metal coordination is converted into DNA release through coupled allosteric remodeling and steric exclusion, establishing a mechanism for signal-dependent transcriptional switching in ArsR/SmtB metalloregulators.

## Introduction

Allosteric transcription factors convert environmental signals into changes in DNA occupancy. For metalloregulators, this requires a physical pathway linking a metal-sensing site to the DNA-binding surface, but complete structural descriptions of the signal-free, DNA-bound, and metal-bound states remain uncommon (1–4).

The ArsR/SmtB family of metalloregulatory proteins comprises a functionally diverse group of prokaryotic transcriptional repressors that respond to a wide range of metal ions, from essential elements such as zinc (Zn^2+^) and nickel to toxic metals like arsenic, lead and cadmium (Cd^2+^). These proteins typically act as homodimers that bind to conserved DNA sequences in the promoter regions of genes involved in metal transport and/or homeostasis (5). Binding of specific metal ions causes protein allosteric conformational changes that reduce their affinity for DNA, leading to the derepression of target gene expression (6–11). Structurally, ArsR/SmtB proteins adopt a conserved winged helix-turn-helix (wHTH) fold, a DNA-binding architecture also found in other metal-responsive transcriptional regulator families such as DtxR/IdeR (12), although coupled to distinct dimerization domains and regulatory mechanisms. ArsR/SmtB proteins coordinate metal ions through cysteine-rich motifs whose number and position vary across homologs and confer distinct metal selectivity (13,14). Although several ArsR/SmtB proteins have been structurally characterized in their apo and metal-bound states, including *Escherichia coli* ArsR (15), *Synechococcus* sp. SmtB (16), *Staphylococcus aureus* CadC (17) and CzrA (18), and *Mycobacterium tuberculosis* CmtR (19) and NmtR (20), the molecular basis of DNA recognition remains poorly understood. Structural insight into DNA-bound ArsR/SmtB proteins has been limited, with previous studies restricted to NMR analysis of the zinc sensor CzrA (21) and the crystal structure of the non-metal-binding regulator NolR (22). A recent structure of the redox-responsive persulfide sensor SqrR in complex with DNA (PDB: 13CT) has further expanded our understanding of DNA recognition within the ArsR superfamily. A direct crystallographic comparison linking cognate-DNA recognition to a metal-bound regulatory state has, to our knowledge, remained lacking for metal-responsive ArsR/SmtB repressors.

*Listeria monocytogenes* (*Lm*) is a major foodborne pathogen and the causative agent of listeriosis. It survives both in the environment as a saprophyte and within mammalian hosts as a facultative intracellular pathogen (23–25), adapting to distinct niches by sensing environmental cues and tightly controlling gene expression (26). The genome of *Lm* EGDe serovar 1/2a includes: *cadA*, which encodes CadA, a Cd^2+^ efflux pump involved in Cd^2+^ resistance (10,27); an upstream *cadC* gene encoding the transcriptional repressor CadC; and between *cadC* and *cadA*, an oppositely oriented gene encoding the lipoprotein signal peptidase LspB (28) (**Figure 1A**). We previously showed that CadC is a Cd^2+^-responsive, sequence-specific DNA-binding regulator that controls the expression of *cadA*, *cadC and lspB* (10). CadC shares 19-46% amino acid sequence identity with other members of the ArsR/SmtB superfamily such as *S. aureus* CadC and CzrA, *Bacillus anthracis* PagR and *Rhodobacter capsulatus* SqrR (**Supplementary Figure S1**). CadC belongs to the family of wHTH-containing transcription factors (InterPro ID: IPR001845). However, unlike *S. aureus* CadC, which harbors two metal-binding sites, including a cysteine-rich α3N site (named after its location at the N-terminus of helix α3) that coordinates thiophilic metals, and a Zn²⁺-specific α5 site located at the dimer interface, *Lm* CadC contains only a single α3N-type site (5,29). Beyond its role in Cd^2+^ detoxification, we also showed that CadC is highly expressed in late infection stages, where it represses genes such as *bsh,* encoding the bile salt hydrolase whose activity becomes detrimental to *Lm* fitness during later stages of infection (10,30). These findings suggest a dual role for CadC in coordinating metal sensing and regulating virulence gene expression during pathogenesis. However, the molecular determinants underlying CadC metal-dependent regulation and DNA-binding specificity are still unknown.

**Figure 1.**
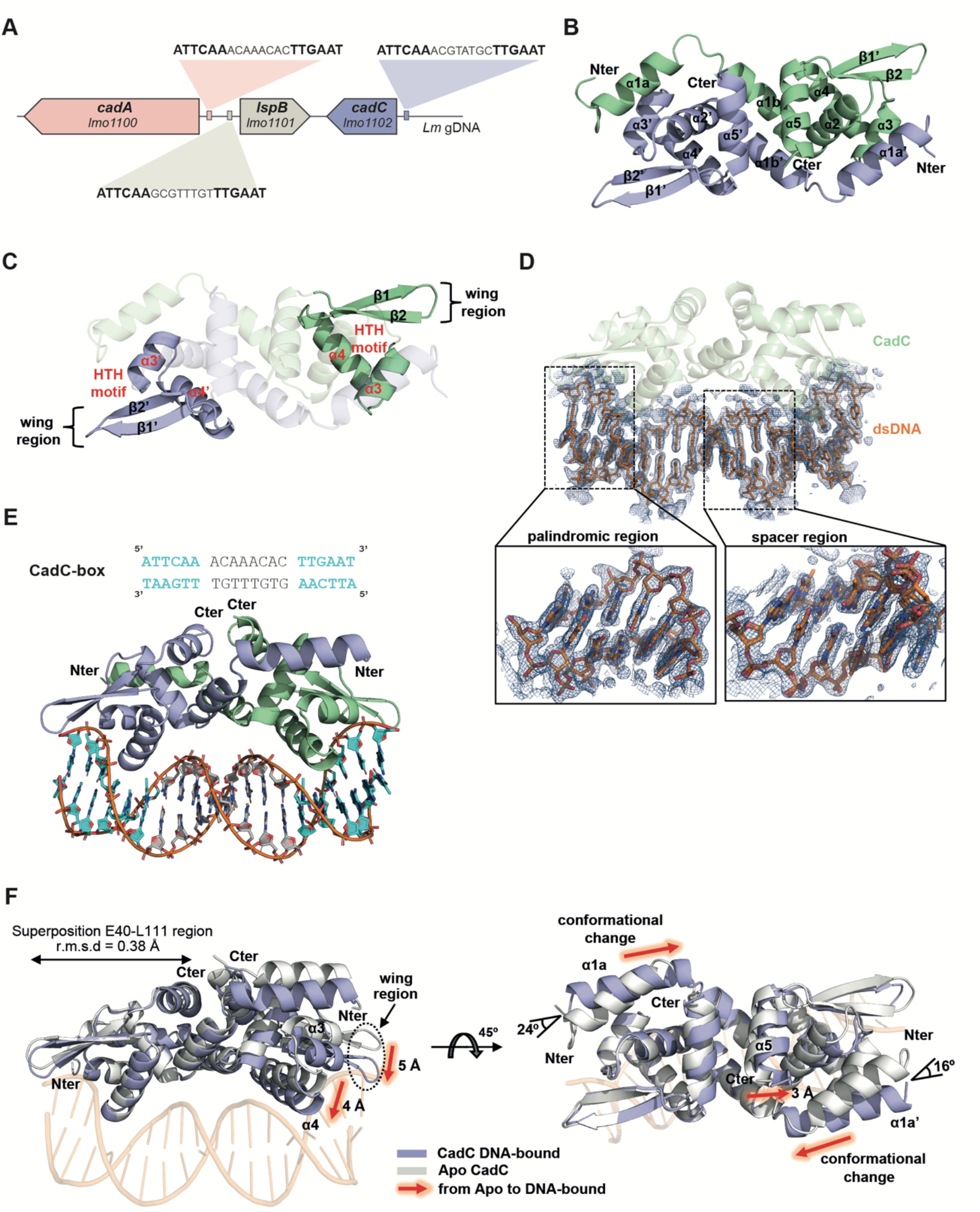
Three-dimensional structure of *Listeria monocytogenes* (*Lm*) CadC. **(A)** Schematic representation of the genomic organization of the *cadA*-*lspB*-*cadC* locus in *Lm*. Genes are coloured in red (*cadA*), gray (*lspB*) and blue (*cadC*), with arrows indicating the direction of transcription. The CadC-binding boxes (CadC-box) located upstream of each gene are shown, with the palindromic regions highlighted in bold. **(B)** Cartoon representation of the CadC dimer with the two monomers coloured in blue and green. The N-(Nter) and C-termini (Cter), as well as secondary structure elements are labelled. **(C)** Cartoon representation highlighting the CadC helix-turn-helix (HTH) motifs formed by helices α3-α4 and the wing region formed by strands β1-β2. **(D)** Electron density map (2Fo-Fc, blue mesh) of the DNA duplex contoured at 1.0σ. The CadC-dsDNA complex is shown with CadC displayed as a pale green cartoon and dsDNA as orange sticks. Lower panels show enlarged views of representative DNA segments, highlighting the continuous electron density surrounding both the phosphate backbone and nucleobases. **(E)** Cartoon representation of the CadC-DNA complex. The nucleotide sequence of the binding site is shown, with the palindromic motif highlighted in cyan in the sequence and structure. **(F)** Superposition of the apo-CadC (white) and DNA-bound CadC (blue) structures through residues 40−111 (Cα RMSD: 0.38 Å). The N-(Nter) and C-termini (Cter) are labelled. Red arrows indicate structural changes from apo to DNA-bound structure.

Here, we combine crystal structures of CadC in its apo, DNA-bound, and Cd²⁺-bound states with mutational, biophysical, transcriptional, and infection analyses. Comparison of these structural states, together with functional analyses, connects operator recognition to metal-induced DNA release and reveals an unexpected methionine-containing α3N coordination site. These data establish a mechanistic framework for metal-gated DNA recognition by CadC and connect this regulatory switch to CadC function during infection.

## Results

### DNA-bound CadC reveals the architecture of operator recognition

To gain insights into the molecular properties of CadC, we determined the crystal structures of CadC (apo-CadC) as well as of CadC in complex with DNA. Data collection and refinement statistics are summarized in **Table 1**.

**Table 1.** Data collection and refinement statistics (molecular replacement). ^a^Values in parentheses are for highest-resolution shell.

| Structure | CadC | CadC<br>Cadmium | CadC<sup>D11-L111</sup><br>DNA (20-mer) |
| --- | --- | --- | --- |
| **Data Collection** |  |  |  |
| Collection date | 2024.09.22 | 2024.09.22 | 2025.02.12 |
| X-ray source | ESRF (ID30b) | ESRF (ID30b) | ESRF (ID30A-3) |
| Detector | Dectris Eiger2_9M | Dectris Eiger2_9M | Dectris Eiger1_4M |
| Space Group | P 3<sub>2</sub> 2 1 | P 2<sub>1</sub> 2<sub>1</sub> 2<sub>1</sub> | C2 |
| Wavelength (Å) | 0,8856 | 0,8856 | 0,9677 |
| **Cell dimensions** |  |  |  |
| a,b,c (Å) | 110, 110, 124 | 63, 65, 103 | 51, 82, 38 |
| α, β, γ (°) | 90, 90, 120 | 90, 90, 90 | 90, 98, 90 |
| Resolution (Å) | 75.5 - 3.0 (3.17 - 3.0) | 40.2 - 2.0 (2.07 - 2.0) | 42.7 - 1.69 (1.8 - 1.69) |
| Number of observations measured | 159 330 (13 388) | 162 817 (15 754) | 79 582 (13 381) |
| Number of unique reflections measured | 17 557 (4 535) | 54 092 (5 376) | 33 103 (5 561) |
| Multiplicity | 4.8 (3.0) | 3.0 (2.9) | 2.4 (2.4) |
| Completeness (%) | 99.7 (98.9) | 98.9 (98.2) | 98.3 (97.7) |
| I/σI | 5.1 (0.96) | 5.8 (0.64) | 8.9 (0.84) |
| Wilson B-factor | 67,7 | 34,8 | 29,4 |
| R<sub>merge</sub> | 0.25 (1.19) | 0.10 (1.73) | 0.05 (1.30) |
| R<sub>pim</sub> | 0.13 (0.79) | 0.07 (1.20) | 0.04 (1.03) |
| CC (1/2) (%) | 99.4 (13.8) | 99.8 (37.3) | 99.8 (56.5) |
| Monomers per asymmetric unit | 6 | 4 | 1 |
| Matthews coefficient (Å<sup>3</sup> Da<sup>-1</sup>) | 2,6 | 1,9 | 1,9 |
| Solvent content (%) | 53,6 | 35,9 | 37,1 |
| **Refinement** |  |  |  |
| R<sub>work</sub>/R<sub>free</sub> (%) | 0.26 / 0.32 | 0.20 / 0.25 | 0.20 / 0.22 |
| Numbers of non-hydrogen atoms | 4 966 | 3 690 | 1 477 |
| Protein | 4 966 | 3 497 | 808 |
| Nucleic acid | - | - | 571 |
| Water | - | 189 | 95 |
| Ligand | - | 4 (cadmium) | 3 (tungstate) |
| **RMSD from standard stereochemistry** |  |  |  |
| Bond lengths (Å) | 0,001 | 0,002 | 0,005 |
| Bond angles (°) | 0,33 | 0,42 | 0,8 |
| **Ramachadran plot statistics** |  |  |  |
| Favored (%) | 97,3 | 99,1 | 99,0 |
| Allowed (%) | 2,7 | 0,9 | 1,0 |
| Disallowed (%) | - | - | - |
| **PDB code** | **9T6R** | **31RT** | **31NU** |

The structure of apo-CadC was determined at 3.0 Å resolution by molecular replacement using a CadC model generated by AlphaFold 3 (31) (**Figure 1B**). The crystal asymmetric unit contains three homodimers (**Supplementary Figure S3**). Pairwise comparison revealed high structural similarity among chains and dimers, with Cα RMSDs ranging between 0.5 and 0.9 Å. Each monomer adopts the canonical topology of the ArsR-SmtB superfamily of transcription factors, namely the wHTH architecture (32) found in many prokaryotic and eukaryotic DNA-binding proteins and known to mediate interactions with DNA. CadC comprises helices α1a, α1b, α2, α3, α4 and α5, together with the antiparallel β-strands β1 and β2. Notably, CadC contains an additional N-terminal helix (α1a) that is absent from many ArsR/SmtB family members, including *S. aureus* CzrA, *R. capsulatus* SqrR and *Synechococcus elongatus* SmtB; consequently, the CadC α1b helix corresponds to the α1 helix typically designated in other ArsR/SmtB regulators. The helix-turn-helix motif is formed by helices α3 and α4 (also known as the recognition helix). Their connecting loop and the “wing” are formed by the antiparallel β-strands β1 and β2 (33) (**Figure 1B and C**). The dimer interface buries approximately 1920 Å^2^ and involves hydrophobic interactions between helices α1b (residues I27, I30, I33, F34, L37) and α5 (residues V102, L105, I106, I109). Additional interprotomer contacts are provided by helix α1a, which extends across the interface and interacts with helix α3 of the opposing subunit (**Figure 1B and Supplementary Figure S4**).

The CadC N-terminal region is disordered in apo-CadC and could not be modelled. We thus reasoned that this may explain why crystals of the full-length protein/DNA complex were not obtained. Thus, to get high-quality crystals of the CadC-DNA complex, we truncated the N-terminal region of CadC, generating CadC^D11-L111^. Co-crystallization was performed using a 20-bp duplex DNA containing the CadC-box of the *cadA* promoter (5′-**ATTCAA**ACAAACAC**TTGAAT**-3′ - in bold are the operator palindromic regions). The crystal structure of the CadC^D11-L111^-DNA complex was solved at 1.7 Å resolution by molecular replacement, using the apo-CadC structure as the search model (**Table 1**).

The refined 2Fo−Fc electron-density map provided continuous density for the DNA duplex, supporting reliable modelling of both the phosphate backbone and the nucleobases **(Figure 1D**). Representative close-up views highlighting different regions of the duplex demonstrate the excellent fit of the refined model to the experimental electron density. In the crystal, two equivalent orientations of the DNA duplex coexist and are related by crystallographic symmetry. As the central region of the operator is non-palindromic, the superposition of these two duplex orientations gives rise to partial occupancies for the corresponding central bases. For clarity, the figures show one of the two equivalent DNA orientations (**Figure 1E**). The CadC dimer engages the operator DNA through its wHTH architecture, with the recognition helix α4 making the major contribution to DNA recognition (**Figure 1E and Supplementary Figure S5**).

Three tungstate ions were identified in the CadC-DNA structure, consistent with the crystallization conditions. All are located outside the CadC-DNA interface and do not participate in DNA recognition (**Supplementary Figure S6**). Comparison of the apo- and DNA-bound structures reveals a more closed relative arrangement of the two protomers within the dimer, while the monomeric core remains highly similar (RMSD = 0.38 Å over 72 Cα atoms) (**Figure 1F**). DNA binding induces localized rearrangements in the wing region and in the recognition helix (α4), which shift by ∼4-5 Å towards the DNA. These movements position key DNA-binding elements for optimal interaction with the duplex (**Figure 1F**, left panel). In addition, the relative orientation of the α1a helix from each monomer differs in the DNA-bound structure as compared to the apo form. Specifically, α1a is rotated by 16° and 24° relative to the apo structure and undergoes a conformational change that extends the helix by approximately one additional turn (**Figure 1F**, right panel).

### Structure-guided mutagenesis defines sequence-specific CadC-DNA recognition

To quantitatively evaluate the importance of the molecular interactions observed in the structure of the CadC-DNA complex, we established a fluorescence anisotropy assay using a 5’-fluorescein-labelled (FAM) 33 bp oligonucleotide encompassing the 20 bp CadC-box of the *cadA* promoter (P*cadA*) in its central region (**Supplementary Table S2**). FAM-DNA titration with increasing concentrations of purified wild-type CadC resulted in an increase in fluorescence anisotropy values, consistent with protein-DNA complex formation (**Figure 2A**, filled circles). The fluorescence anisotropy titration data were fitted with a Hill equation, yielding an apparent half-saturation concentration (K_1/2_) of 0.38 ± 0.05 µM and a Hill coefficient of 3.84 ± 1.11 (**Figure 2A**, filled circles).

**Figure 2.**
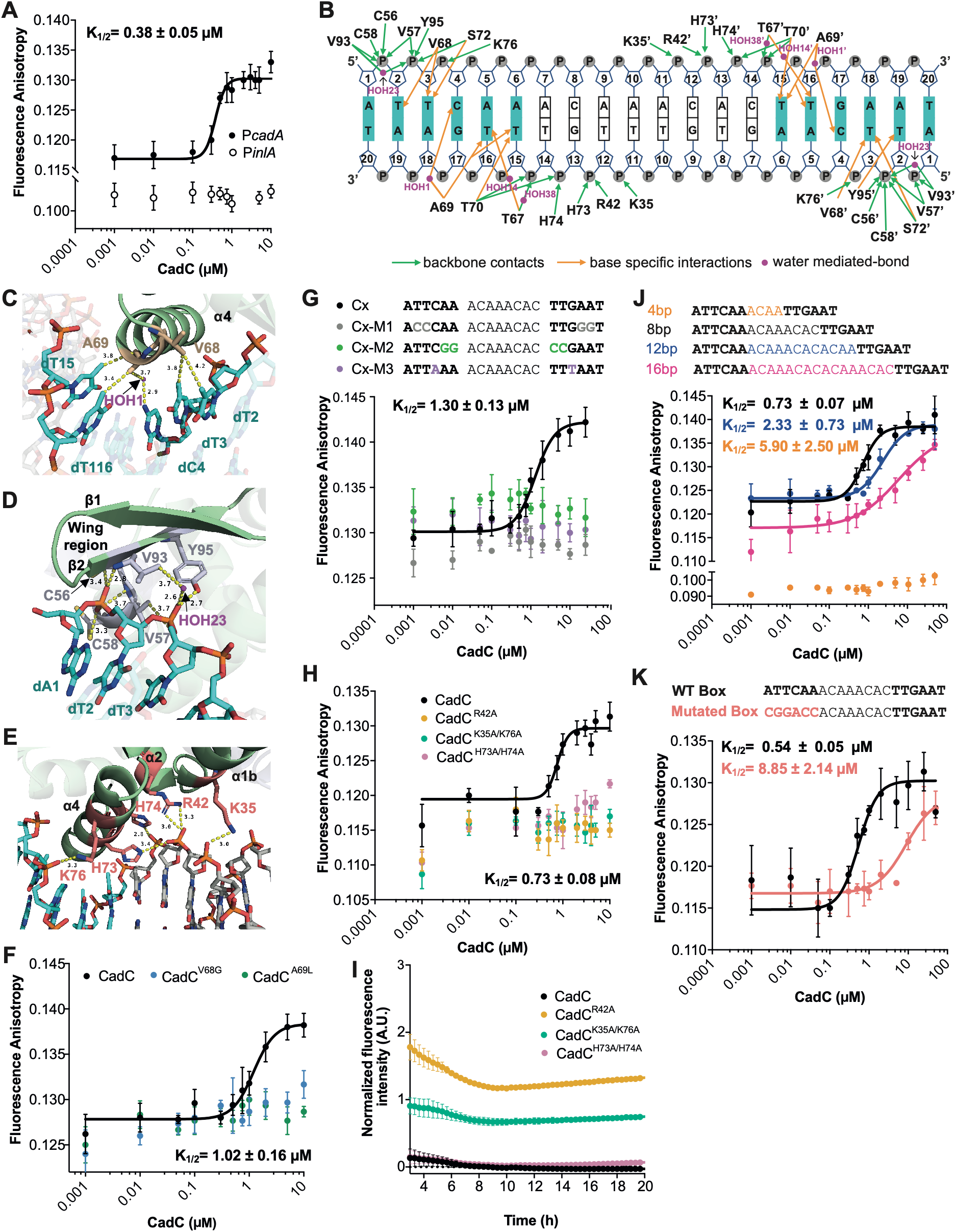
CadC-DNA interactions. **(A)** Fluorescence anisotropy analysis of CadC binding to the *cadA* promoter (P*cadA*) CadC-Box. 5′-fluorescein (FAM)-labeled DNA was titrated with increasing concentrations of purified CadC. The negative control corresponds to the unrelated P*inlA* DNA sequence**. (B)** Schematic representation of the interactions between CadC^D11-L111^ and the CadC-box. Nucleotides are shown as rectangles with one-letter codes, while phosphate and ribose groups are depicted as circles and pentagons, respectively. Residues from chain B of the CadC homodimer are indicated with an apostrophe following the amino acid number. Green arrows denote backbone contacts, orange arrows denote base-specific interactions, and magenta circles indicate water-mediated hydrogen bonds. The consensus target DNA sequence is highlighted in cyan. Water molecules are shown as magenta circles, labeled with their PDB identifiers. **(C)** Close-up view of base-specific interactions involving residues V68 and A69 in helix α4. Nucleotides from the 5′ and 3′ strands are coloured in cyan and labelled. Residues and water molecules involved in the interactions are shown in wheat and magenta, respectively. Dashed lines indicate the contact distances in angstroms. **(D)** Close-up view of the interactions between one CadC^D11-L111^ monomer and the palindromic region in the wing domain, as in (C). **(E)** Representation of the backbone interactions between one CadC^D11-L111^ monomer and the CadC-box, as in (C). Fluorescence anisotropy analysis of: **(F)** CadC (black circles) and the mutants CadC^V68G^ (blue circles) and CadC^A69L^ (green circles) binding to the CadC-box; **(G)** CadC binding to DNA containing mutations in the CadC-box; and **(H)** CadC (black circles) and the mutants CadC^R42A^ (yellow circles), CadC^K35A/K76A^ (green circles) and CadC^H73A/H74A^ (pink circles) binding to the CadC-box. **(I)** Normalized GFP fluorescence in *Lm* strains carrying a P*cadA*-GFP transcriptional reporter fusion under the control of CadC mutants R42A, K35A/K76A, H73A/H74A. Bacteria were grown in a chemically defined medium at 37 °C with agitation and GFP fluorescence (excitation 470/15 nm) was monitored every 20 min. Fluorescence anisotropy of: **(J)** CadC binding to CadC-boxes with different spacer lengths (4, 12 or 16 bp), compared with the canonical spacer sequence (8 bp); and to **(K)** the CadC-box with one half-site mutated. For **(F)**, **(G)**, **(H)**, **(I)**, **(J)** and **(K)**, 5’FAM-labelled oligonucleotides were titrated with increasing concentrations (0-10 µM) of CadC or mutant proteins at 22 °C. Data represent the mean ± SD of three independent experiments. Data were fitted with the Hill equation. Fitted parameters are shown in **Supplementary Table S3.**

In contrast, when using a control oligonucleotide derived from an unrelated gene promoter lacking a CadC-box (P*inlA*), the fluorescence anisotropy remained unchanged (**Figure 2A**, open circles), indicating the absence of binding and confirming the specificity of the CadC-CadC-box interaction. Size-exclusion chromatography further supported the formation of a complex between CadC and the DNA oligonucleotide containing the CadC-box from P*cadA*, with the complex eluting as a single peak at 13.7 ml, earlier than CadC alone (15.8 ml) or free DNA (14.1 ml) (**Figure 3D**). SDS-PAGE and fluorescence measurements confirmed the presence of both CadC and fluorescein-labeled DNA in the complex elution peak (**Figure 3D**). Comparable DNA binding and affinities were obtained for CadC binding to the CadC boxes present in the *lspB* (K_1/2_ = 0.52 ± 0.07 µM) and *cadC* promoters (K_1/2_ = 0.61 ± 0.05 µM) (**Supplementary Figure S7A and B,** respectively). These interactions were further confirmed by size-exclusion chromatography (**Supplementary Figure S7C**). It is worth noting that, across seven independent fluorescence anisotropy experiments, the apparent K_1/2_ for CadC binding to DNA, ranged from 0.38 to 1.30 µM (overall mean ± s.d., 0.74 ± 0.32 µM). To account for inter-assay variability, wild-type CadC was included in all assay plates and mutant proteins were evaluated relative to that specific wild-type control. In addition, in many of the anisotropy experiments, fitting the data with the Hill equation yielded Hill coefficients greater than the expected number of binding sites (two) on the DNA (see below and **Supplementary Table 3**). The molecular basis for this effect is unclear and, therefore, we compare only K_1/2_ values across experiments.

**Figure 3.**
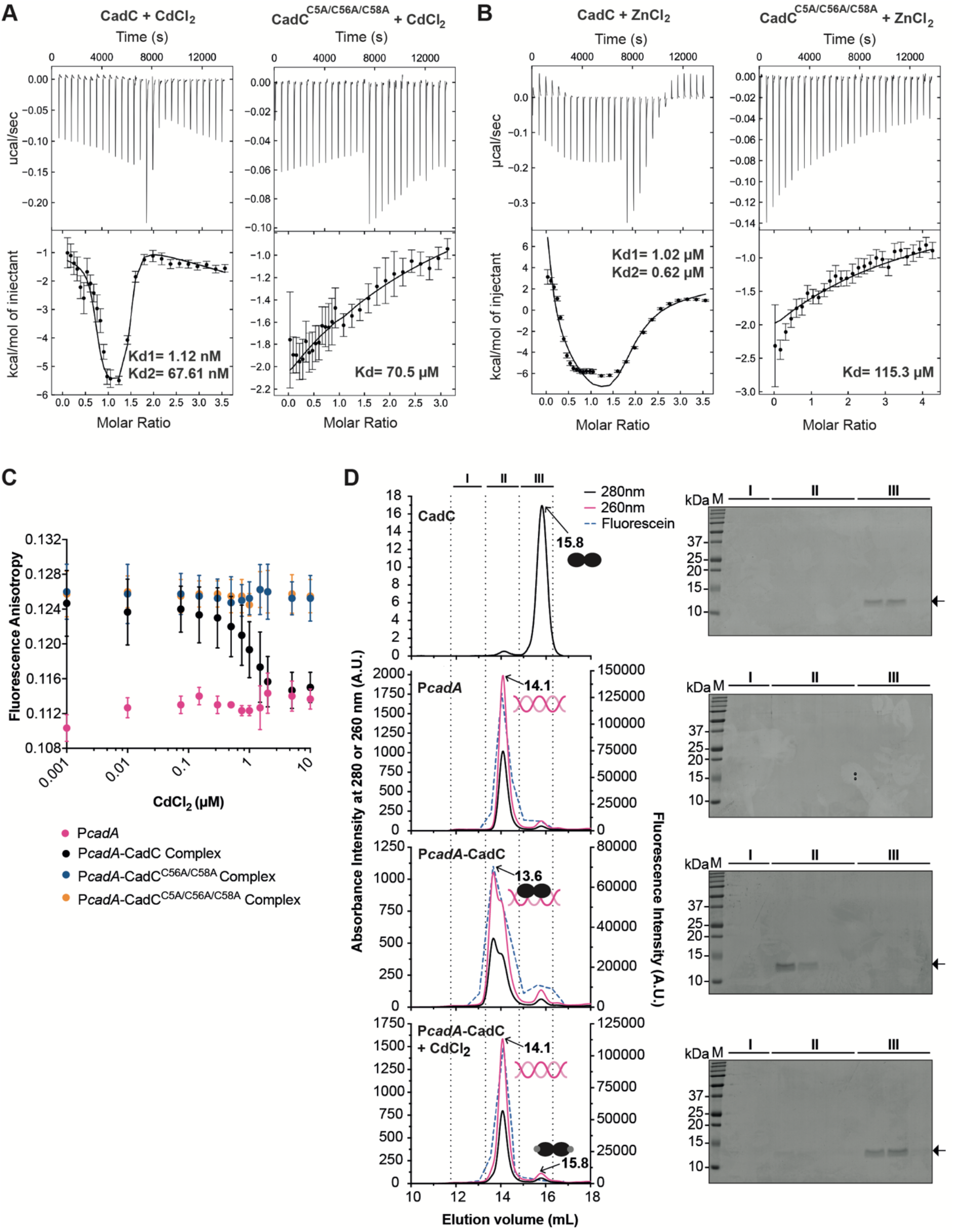
CadC binds cadmium in a two-event binding process. **(A-B)** ITC measurements of the interaction of CadC (10 µM) with divalent metals (200 µM): **(A)** CdCl_2_ and **(B)** ZnCl_2_ titrated into wild-type CadC or CadC^C5A/C56A/C58A^ at 15 °C. Upper panels show raw data plotted as power over time, and the lower panels show heat released when metal binds to CadC as a function of relative concentrations. The heat of dilution was corrected by subtracting the baseline obtained from the control titration (200 µM CdCl_2_ into 50 mM HEPES pH 7.5, 200 mM NaCl, 1 mM TCEP) from the raw data. Solid lines correspond to the best fit to a two-site binding model with two symmetry-related sites and macroscopic binding constants. Data shown are representative of two independent experiments yielding similar results (compiled in **Table 2**). **(C)** Fluorescence anisotropy analysis of P*cadA* in complex with either wild-type CadC (P*cadA*-CadC) or mutated in cysteine residues (P*cadA*-CadC^C56A/C58A^ and P*cadA*-CadC^C5A/C56A/C58A^), titrated with increasing concentrations of CdCl_2_. P*cadA* alone (pink circles dots) was included as negative control. Data represent the mean ± SD of three independent experiments and were fitted with a competition model in DynaFit. Fitted parameters are shown in **Supplementary Table S4. (D)** Cadmium disrupts CadC binding to the CadC box within the *cadA* promoter. Size exclusion chromatography profiles were monitored at 280 nm (CadC - black line) and 260 nm (P*cadA* - pink line). FAM fluorescence (blue dashed line) was also measured for each collected fraction in a fluorescence plate reader. Chromatograms shown correspond to samples containing: CadC; FAM-P*cadA* alone, FAM-P*cadA* complexed with CadC in the absence of CdCl_2_, and CadC displaced from FAM-P*cadA* in the presence of CdCl_2_. Elution times for each peak are indicated above the chromatograms. Eluted fractions were labelled as I, II and III, and were analysed by SDS-PAGE electrophoresis, where the protein bands correspond to the monomeric CadC. Lane M, molecular mass markers (kDa; sizes shown on the left).

**Table 2.**
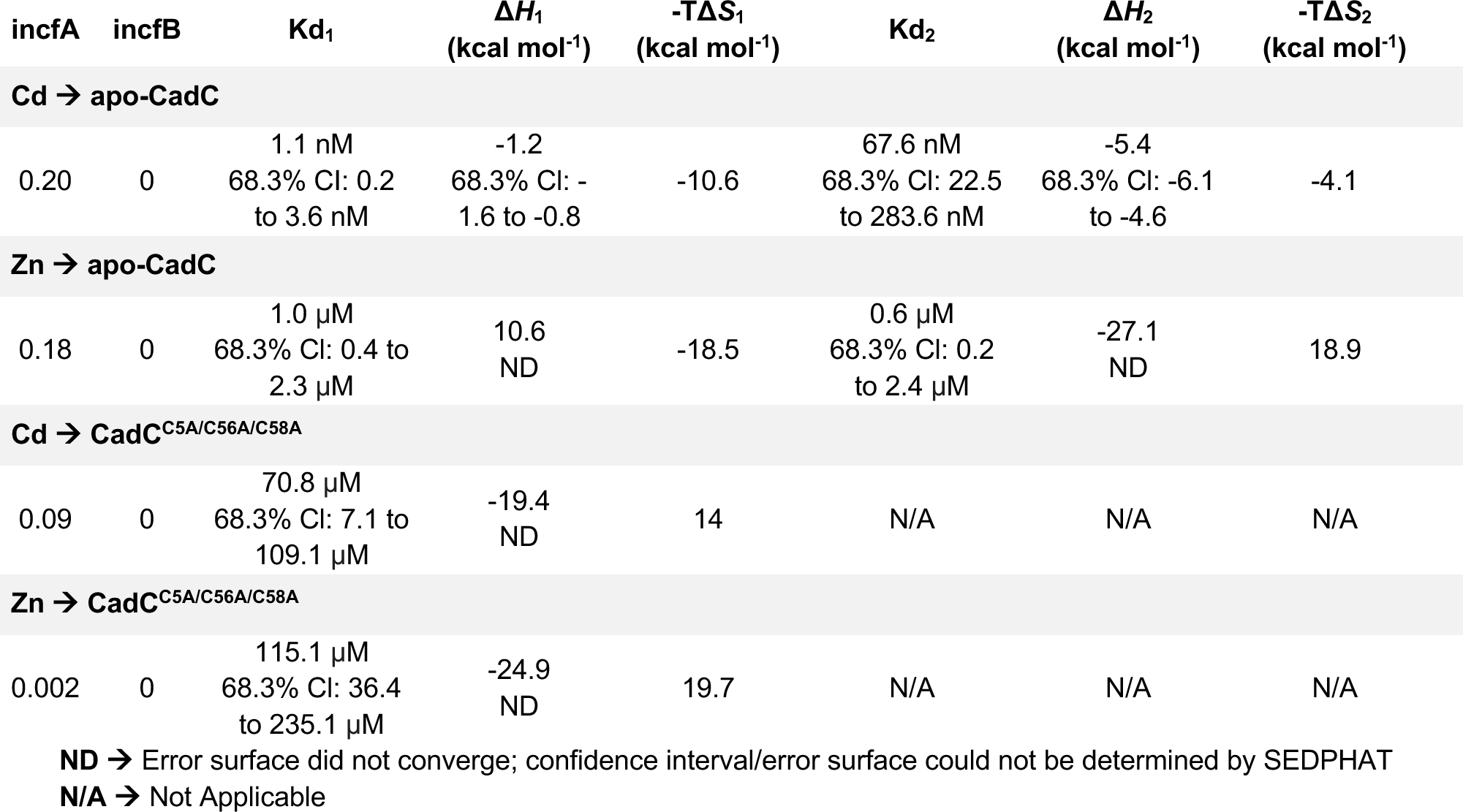
Thermodynamic parameters derived from direct analysis of isothermal titration calorimetry (ITC) experiments.

PDBePISA analysis (34) of the CadC-DNA complex identified 30 residues of the CadC dimer involved in 48 protein-DNA contacts, with each monomer contributing nearly identical interactions with one half-site of the palindromic operator (**Figure 2B-E**). In the HTH motifs, the recognition α-helices (α4/α4′) insert into two consecutive major grooves of the operator DNA (**Supplementary Figure S5A**) and establish base-specific contacts through residues V68, A69, T70 and S72 together with T67 in the turn region of the HTH motif (**Supplementary Figure S5B**). V68 and A69 stand out, since they only interact with the nucleotide bases while the other residues also interact with the DNA backbone (**Figure 2C**). The sidechain of V68 forms van der Waals interactions with the methyl groups of two thymine bases (T2 and T3) within the palindromic sequence. Alanine 69 interacts with three bases through a combination of direct and water-mediated contacts, including a water-mediated interaction between its methyl group and the N4 atom of cytosine 4, as well as two van der Waals contacts with the methyl groups of thymine bases T15 and T16. We assessed the contribution of V68 and A69 to DNA binding by fluorescence anisotropy (**Figure 2F)**.

In contrast to wild-type CadC, substitutions V68G or A69L severely impaired DNA binding, demonstrating that V68 and A69 are determinant for CadC recognition of the cognate DNA operator.

We also dissected the role of the DNA bases in the interactions with CadC. DNA variants containing point mutations within the palindrome sequence severely disrupted complex formation, demonstrating the strict sequence specificity of CadC-DNA recognition and the specific importance of the contacts established between the protein and the thymine bases (**Figure 2G**).

In the wing region (**Figure 2B and D**), V93 interacts with the phosphate groups of thymine 2 and 3. In addition, Y95, highly conserved not only among ArsR/SmtB family members but also in the YodB/QsrR family of regulators **(Supplementary Figure S1)**, establishes a direct hydrogen bond between its side-chain hydroxyl group and the phosphate of thymine 3. Notably, an analogous tyrosine-DNA phosphate interaction has also been described for QsrR (35), highlighting a conserved role for this residue in stabilizing protein-DNA complexes across related transcriptional regulator families. Finally, a cluster of residues, including K35 and the highly conserved R42, H73, H74 and K76, form an extensive network of electrostatic and hydrogen-bonding interactions with the phosphate backbone of both DNA strands (**Figure 2B and E**). To assess the functional relevance of these contacts, we performed fluorescence anisotropy assay with the mutants CadC^R42A^; CadC^K35A/K76A^ and CadC^H73A/H74A^ (**Figure 2H**). Mutation of R42 alone, or simultaneous substitution of K35/K76 or H73/H74 abolished detectable CadC interaction with DNA, highlighting the essential role of these residues in CadC-DNA recognition (**Figure 2H**). All CadC variants displayed thermal stability well above room temperature (**Supplementary Figure S8A**), excluding protein misfolding as an explanation for these effects.

To determine the role of these residues *in vivo*, we constructed a *Lm* Δ*cadC* strain constitutively expressing mCherry and carrying a chromosome-integrated reporter plasmid that drives GFP expression under the control of the *cadA* promoter (*Lm* Δ*cadC* P*cadA*), thus enabling simultaneous visualization of bacteria (red) and promoter activity (green). Wild-type CadC and CadC mutants (CadC^R42A^; CadC^K35A/K76A^ and CadC^H73A/H74A^) were expressed in the *Lm* Δ*cadC* P*cadA* strain from a non-integrative plasmid. *The Lm* Δ*cadC* P*cadA* strain expressing wild-type CadC or CadC^H73A/H74A^ variant barely expressed GFP, indicating a repressed expression of *cadA.* In contrast, strains expressing CadC^R42A^ or CadC^K35A/K76A^ mutants displayed constitutive GFP expression (**Figure 2I**) comparable to that observed in the Δ*cadC* deletion mutant (**Supplementary Figure S9**), showing that *cadA* expression was likely due to loss of CadC-mediated repression. These data indicate that the H73A/H74A mutation strongly compromises DNA binding *in vitro* but does not abolish CadC-mediated repression under the specific *in vivo* reporter conditions used. Furthermore, consistent with the *in vitro* findings, *in vivo* GFP reporter assays corroborated the functional relevance of the R42 and K35/K76 residues. Importantly, protein levels of these CadC mutants were comparable to those of wild-type CadC in the presence of cadmium (**Supplementary Figure S8B**), and none of the mutations affected bacterial growth (**Supplementary Figure S10**).

We next tested how the spacing between the palindromic half-sites of the CadC-box influences binding. Shortening the canonical 8-bp spacer to 4 bp abolished detectable binding, whereas increasing it to 12 bp reduced affinity by ∼3-fold (apparent K_1/2_ = 2.33 ± 0.73 µM) and increasing it to 16 bp reduced affinity by ∼8-fold (apparent K_1/2_ = 5.92 ± 2.50 µM) (**Figure 2J**). To test whether the longer-spacer complexes could reflect engagement of only one half-site, we introduced a previously characterized binding-defective substitution (10) into the first inverted repeat of the CadC-box of the P*cadA* sequence. Binding was strongly impaired (apparent K_1/2_ = 8.85 ± 2.14 µM), a ∼16-fold reduction relative to the wild-type operator (**Figure 2K**). The similar apparent K_1/2_ values for the single-repeat and 16-bp-spacer variants are consistent with predominantly single-half-site engagement in the latter. Thus, the canonical operator geometry is strongly favored for high-affinity binding, although extension of the spacer does not abolish CadC-DNA association.

Altogether, these findings indicate that optimal CadC binding to DNA requires a precise inverted repeat spacing for stable DNA engagement, likely reflecting the molecular dimensions of the protein and the relative positioning of the major grooves in B-DNA, and highlighting a potential constraint on operator architecture shared by related dimeric regulators.

### Cadmium binding disrupts CadC-DNA complexes through the α3N sensor

A defining feature of ArsR/SmtB family regulators is their ability to sense metal ions to control gene transcription. Because CadC is a Cd²⁺-responsive transcriptional regulator that controls the expression of *cadA, lspB*, and *cadC* in *Lm* (Figure 1A) (10,22), we used isothermal titration calorimetry (ITC) to characterize Cd²⁺ binding. Titration of CdCl₂ into CadC revealed two binding events (Figure 3A). Fitting the data with a two-symmetric-site model using macroscopic binding constants was consistent with a very high-affinity first Cd^2+^-binding event, with an apparent Kd in the low-nanomolar range, followed by a lower-affinity event (best-fit Kd values, 1.12 and 67.61 nM, respectively) (**Table 2**). Projection-error analysis yielded bounded 68.3% confidence intervals for the fitted affinity parameters (**Supplementary Figure S11**), indicating that the solution is well constrained within the adopted model. However, the first event has a C value (C= [Protein]/Kd) far above the range optimally resolved by direct ITC under our experimental conditions. We therefore interpret the absolute magnitude of this best-fit affinity cautiously and refer to it as a very-high-affinity event rather than as a precisely determined nanomolar Kd. Simulations in **Supplementary Figure S12** show that the lower-affinity event can broaden the titration profile in a manner analogous to an intrinsic competitor. These simulations support the plausibility of the two-event model but do not independently establish the absolute affinity of the first event. Within the adopted two-event model, the large separation between the fitted macroscopic constants is consistent with apparent negative cooperativity, although alternative models cannot be formally excluded. The second event was also more exothermic than the first (-5.4 versus -1.2 kcal mol⁻¹; **Table 2**). A buffer control showed only dilution heat (**Figure 3A**, middle panel). CadC is predicted to contain a single α3N metal-sensing site formed by residues C5, C9, C56, and C58 (25,35). Consistent with this assignment, Cd²⁺ titration of the C5A/C56A/C58A mutant produced a qualitatively different profile that was fitted by a single low-affinity event (apparent Kd ≈ 70.8 µM; **Figure 3A**, right panel), supporting an essential contribution of the α3N cysteines to Cd²⁺ coordination.

To assess metal selectivity, we tested two physiologically relevant divalent cations: manganese (Mn^2+^), an essential cofactor in numerous biological processes, and zinc (Zn^2+^), the second most abundant trace element in mammalian cells after iron and an essential micronutrient in human physiology (36,37). Mn^2+^ showed no detectable interaction with CadC (**Supplementary Figure S13A**). In contrast, Zn^2+^ also bound CadC *via* two fitted binding events, with apparent affinities in the micromolar range (Kd1 ≈ 1.0 µM; Kd2 ≈ 0.6 µM) and substantially weaker binding than observed for Cd²⁺ **(Table 2)**. As observed for Cd^2+^, the second Zn^2+^ binding event was associated with a substantially larger enthalpy change (−27.1 vs 10.6 kcal mol⁻¹; **Figure 3B**, left panel; **Table 2**). Consistent with what was observed with CdCl_2_, titration of ZnCl_2_ into the triple cysteine mutant (CadC^C5A/C56A/C58A^) yielded a single, weaker binding event (Kd ≈ 115.1 µM), further supporting the involvement of C5, C56, and C58 in metal coordination (**Figure 3B**, right panel). Overall, these data show that, under the conditions tested, CadC binds Cd^2+^ and Zn^2+^, although it binds Cd^2+^ with substantially higher affinity.

Metal binding is expected to regulate the interaction between CadC and its cognate DNA. We therefore analyzed the effect of Cd^2+^ on preformed CadC-DNA complexes using fluorescence anisotropy. Titration with increasing CdCl_2_ concentrations resulted in a dose-dependent decrease of fluorescence anisotropy (**Figure 3C**), an effect not observed with MnCl_2_ (**Supplementary Figure S13B**), consistent with Cd^2+^-induced dissociation of CadC from the CadC-box (P*cadA*). A similar effect was observed using the P*cadC* DNA (**Supplementary Figure S14A**). By contrast, titration with increasing ZnCl_2_ concentrations (up to 10 µM) had no detectable effect on CadC-DNA binding (**Supplementary Figure S14B**). Thus, Zn²⁺ did not measurably dissociate the purified complex within the experimentally accessible concentration range; higher concentrations could not be tested because they caused protein precipitation. Size-exclusion chromatography independently supported Cd^2+^-dependent disruption of the CadC-DNA complex (**Figure 3D**). After addition of 25 µM Cd²⁺, CadC and DNA separated into distinct fractions and the DNA eluted at the position of free DNA (14.1 ml). Altogether, these results demonstrate that Cd^2+^ binds CadC with high affinity and efficiently disrupts CadC-DNA complex *in vitro*. To confirm that cysteines in the α3N metal-binding site are involved in the regulation of DNA binding, we made use of the CadC triple cysteine mutant (CadC^C5A/C56A/C58A^) and generated an additional double mutant (CadC^C56A/C58A^). These mutations did not affect protein stability (**Supplementary Figure S8A**) and mutant proteins still bind DNA with similar affinity to wild-type CadC (**Supplementary Figure S15**). Unlike the complex with wild-type protein under the same conditions (FAM-P*cadA*-CadC), titration of pre-formed FAM-P*cadA*-CadC^C56A/C58A^ and FAM-P*cadA*-CadC^C5A/C56A/C58A^ complexes with increasing concentrations of CdCl_2_ did not result in fluorescence anisotropy changes, (**Figure 3C**), confirming the critical role of cysteines located at the metal binding site in the Cd^2+^-dependent regulation of the CadC-DNA interaction.

To determine the impact of the cation metals *in vivo*, we constructed a *Lm* strain constitutively expressing mCherry and carrying a GFP transcriptional reporter under the control of the *cadA* promoter (P*cadA*). Mid-log phase bacteria were diluted (1:20) and, after 5 h of growth, increasing concentrations of CdCl_2_ (1, 5, 10, 25 and 50 µM) were added. Addition of CdCl_2_ triggered a rapid, CdCl_2_ concentration-dependent increase in GFP fluorescence, indicating derepression of *cadA* expression (**Figure 4A**). Importantly, CdCl_2_ did not affect bacterial growth under the conditions tested (**Supplementary Figure S16A**). A comparable response was observed with a P*cadC*-GFP reporter strain (**Supplementary Figure S16B and C**). In parallel, we examined the regulatory effect of Zn^2+^. Addition of ZnCl_2_ after 5 h of growth also triggered an increase in GFP fluorescence (**Figure 4B**), indicating derepression of *cadA*. The response was weaker than that observed with CdCl_2_, consistent with the lower affinity of CadC for Zn^2+^. A P*cadC* transcriptional reporter also showed ZnCl_2_-dependent derepression (**Supplementary Figure S17A**) and growth curves confirmed that ZnCl_2_ did not impair bacterial growth (**Supplementary Figure S17B and C**). We observed an increase in GFP signal when bacteria entered the stationary growth phase (at approximately 9-10 h under our experimental conditions), even in the absence of Cd²⁺. The basis for this late increase is currently unclear and may reflect growth phase-dependent changes in reporter fluorescence, promoter activity or cellular physiology. However, this does not affect the interpretation of the Cd²⁺-dependent response, because our analysis focuses on earlier time points, before the onset of this stationary-phase-associated GFP increase.

**Figure 4.**
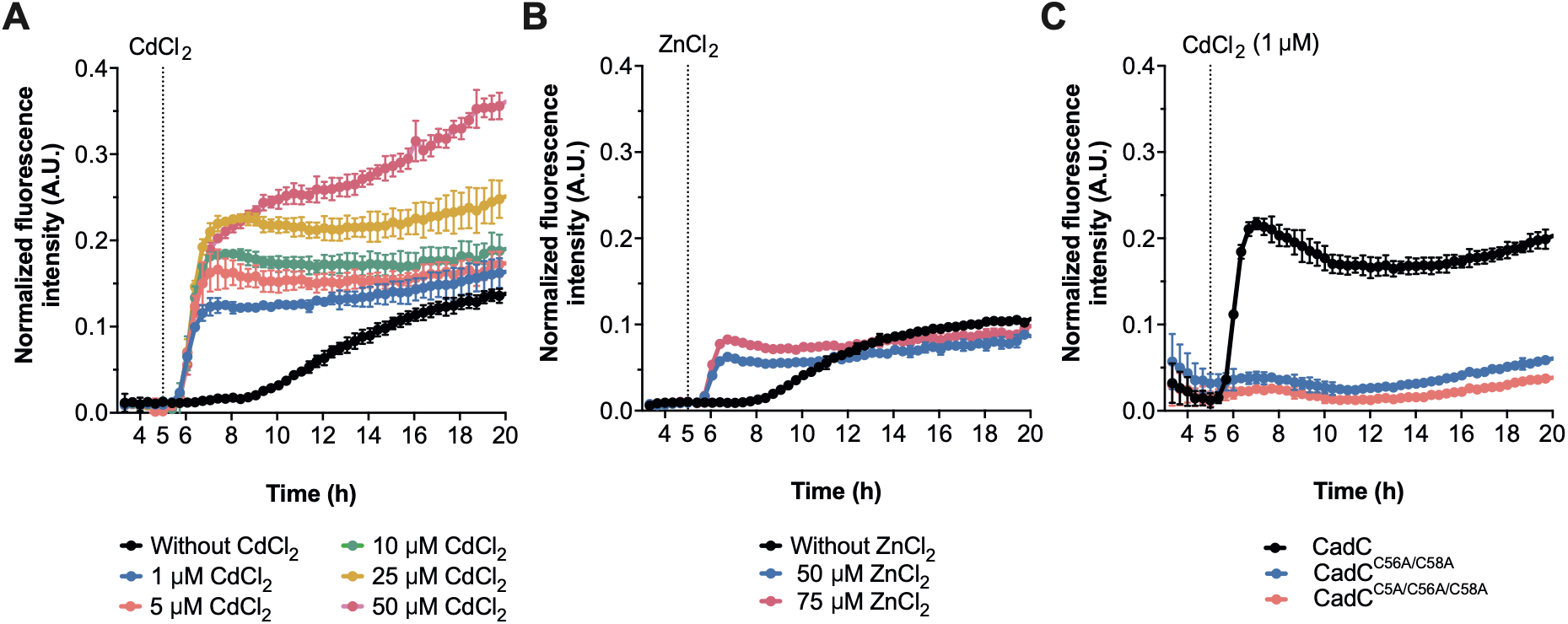
Cadmium and zinc modulate CadC-dependent transcription *in vivo*. Normalized GFP fluorescence of *Lm* strains constitutively expressing mCherry and carrying the P*cadA*-GFP transcriptional reporter fusion. **(A)** CdCl2 (1-50 µM) or **(B)** ZnCl2 (50 or 75 µM) was added at 5 h of growth (dashed line). **(C)** Normalized GFP fluorescence of mCherry *Lm* Δ*cadC* expressing the P*cadA*-GFP transcriptional reporter fusion and CadC cysteine mutants (CadC^C56A/C58A^ and CadC^C5A/C56A/C58A^). CdCl2 was added to a final concentration of 1 µM after 5 h of growth (dashed line). For all experiments, GFP fluorescence (excitation 470/15 nm) was recorded every 20 min and normalized to mCherry fluorescence. Data represent the mean ± SD of three independent experiments.

We next tested whether the cadmium-dependent transcriptional response was mediated by the α3N metal-binding site. We constructed a reporter strain where *Lm* Δ*cadC* P*cadA* was transformed with a plasmid expressing wild-type CadC or CadC cysteine mutants (CadC^C56A/C58A^ and CadC^C5A/C56A/C58A^). While in the presence of wild-type CadC, GFP expression was detected upon addition of CdCl_2_, in both cysteine mutants CdCl_2_-induced derepression was abolished (**Figure 4C**). Levels of CadC cysteine mutants were comparable to those of wild-type CadC (**Supplementary Figure S8B**) and bacterial growth was similar for all CadC variants, even in the presence of CdCl_2_ (**Supplementary Figure S18**). Altogether, these results show that the α3N metal-binding site is required for cadmium-dependent regulation of CadC *in vivo*, and that Cd²⁺ is a more potent modulator of CadC activity than Zn²⁺ under the conditions tested.

### Cadmium stabilizes a DNA-incompatible CadC conformation

To understand the molecular details of the interaction between Cd^2+^ and CadC, and how this regulates DNA binding, we determined the structure of the complex formed by full-length CadC and the metal cation at 2.0 Å resolution (data collection and refinement statistics are summarized in **Table 1**). The crystallographic asymmetric unit contains two homodimers with each protein subunit coordinating a single cadmium ion. The final crystallographic model of the complex (**Figure 5A and B**) lacks only the first residue at the N-termini, which is not defined in the electron density map. The metal at this site was assigned as Cd²⁺ based on the crystallization conditions, the position and coordination geometry of the metal-associated electron density (**Figure 5C**), comparison with the apo-CadC structure, and the mutational and biochemical data supporting the involvement of the α3N site in Cd²⁺-dependent regulation.

**Figure 5.**
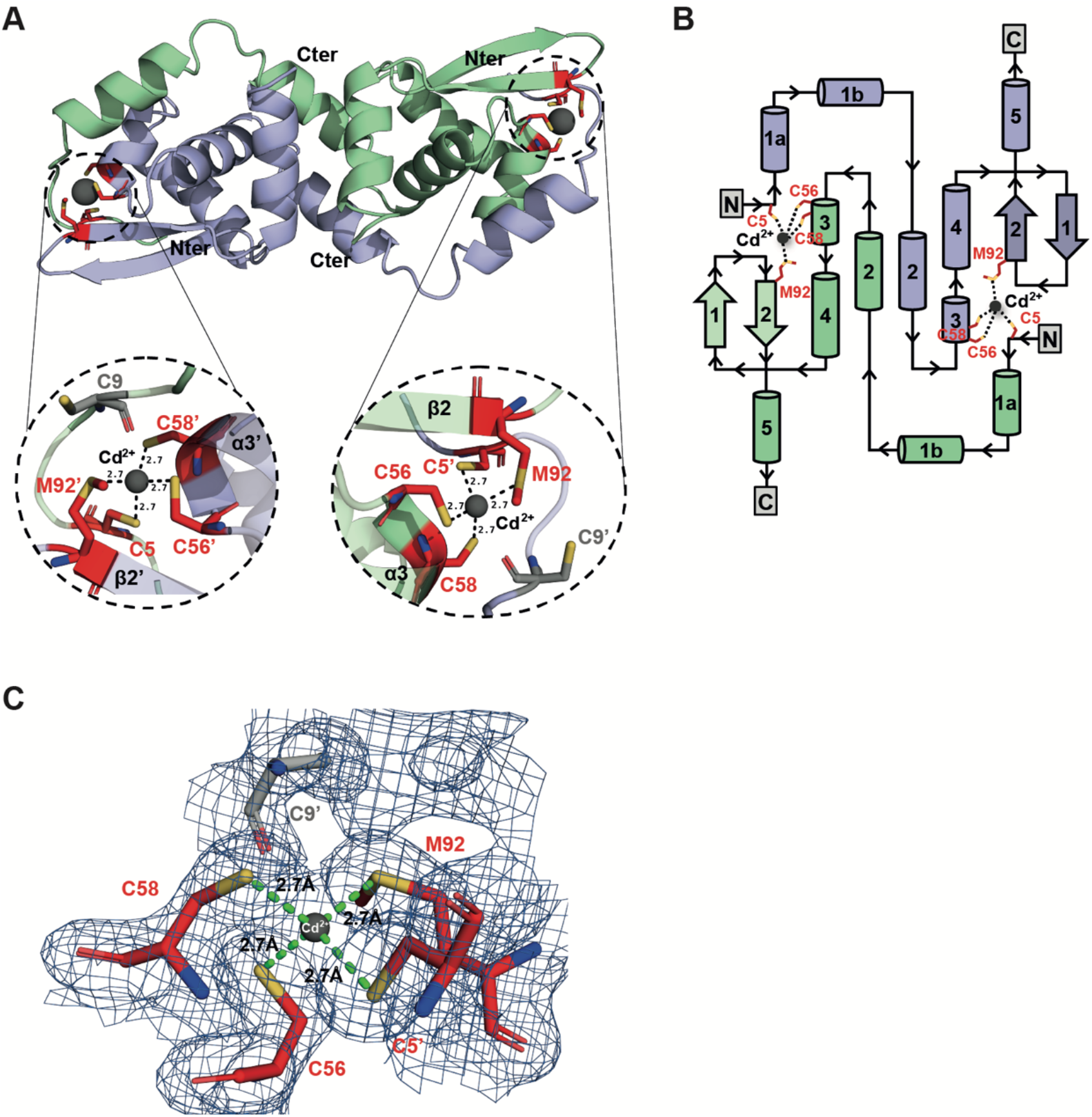
Cadmium-bound CadC structure reveals a distinct metal-binding geometry. **(A)** Cartoon representation of the CadC dimer bound to cadmium. The N-(Nter) and C-termini (Cter) are labeled. The insets show a close-up of the active sites, including the cadmium ion (Cd^2+^; grey sphere) and the coordinating residues from CadC (red). Dashed lines indicate distances between cadmium and amino acid residues in angstroms. **(B)** Topological representation of CadC with monomers colored as in Figure 1B. Secondary-structure elements are depicted and labelled. Cd binding sites and CadC residues involved in its coordination (C5, C56, C58 and M92) are also indicated. **(C)** Electron density map of the Cd^2+^-binding site. The Cd^2+^-binding site is shown with the coordinating residues from CadC (red sticks) and Cd^2+^ ion (grey sphere) and enveloped by the corresponding 2Fo-Fc electron density map contoured at 1.0 σ (blue mesh). Green dashed lines indicate metal-ligand coordination distances (Å).

Analysis of the metal-binding site revealed that Cd^2+^ is coordinated by C5, C56, and C58 (**Figure 5A and C**), consistent with our biochemical measurements (**Figure 3C),** and the canonical α3N arrangement observed in other ArsR/SmtB regulators. However, in contrast to *S. aureus* CadC and related homologs, where Cd^2+^ coordination typically involves a fourth cysteine residue (e.g., C11 in *S. aureus* CadC, corresponding to the conserved C9 in CadC), the Cd^2+^-bound CadC structure shows that C9 does not participate in metal coordination. Instead, the thioether group in the side-chain of M92 provides a sulphur donor to complete the coordination (**Figure 5A-C**). Thus, despite retaining the conserved residue that often contributes to Cd^2+^ coordination in α3N-site ArsR/SmtB regulators, CadC adopts a distinct metal-binding geometry in which M92, rather than C9, completes the observed coordination environment.

A structural superposition of the DNA-CadC complex with the Cd^2+^-CadC structure through one of the subunits reveals that several local conformational differences affect regions involved in DNA recognition (**Figure 6**). In the DNA-bound structure, the wing region and the HTH motif (α3 and α4) are positioned to engage the DNA duplex, whereas in the Cd^2+^-bound structure these elements are shifted by ∼8 Å away from the DNA-binding surface. Consistently, residues implicated in DNA interaction, including V68, A69, H73, H74 and K76 (**Figure 2F, H and I**) are positioned farther from the DNA backbone (**Figure 6**). Moreover, cadmium binding induces a positional shift of ∼3 Å in R42 from helix α2, a key residue involved in DNA binding (**Figure 2H** and **I**), further repositioning it away from the DNA backbone (**Figure 6**). A major structural difference is also observed at the N-terminal region: in the Cd^2+^-bound structure, helix α1a undergoes partial unwinding and reorientation (∼25°) relative to the DNA-bound conformation, altering the positioning of the N-terminal segment (**Figure 6**).

**Figure 6.**
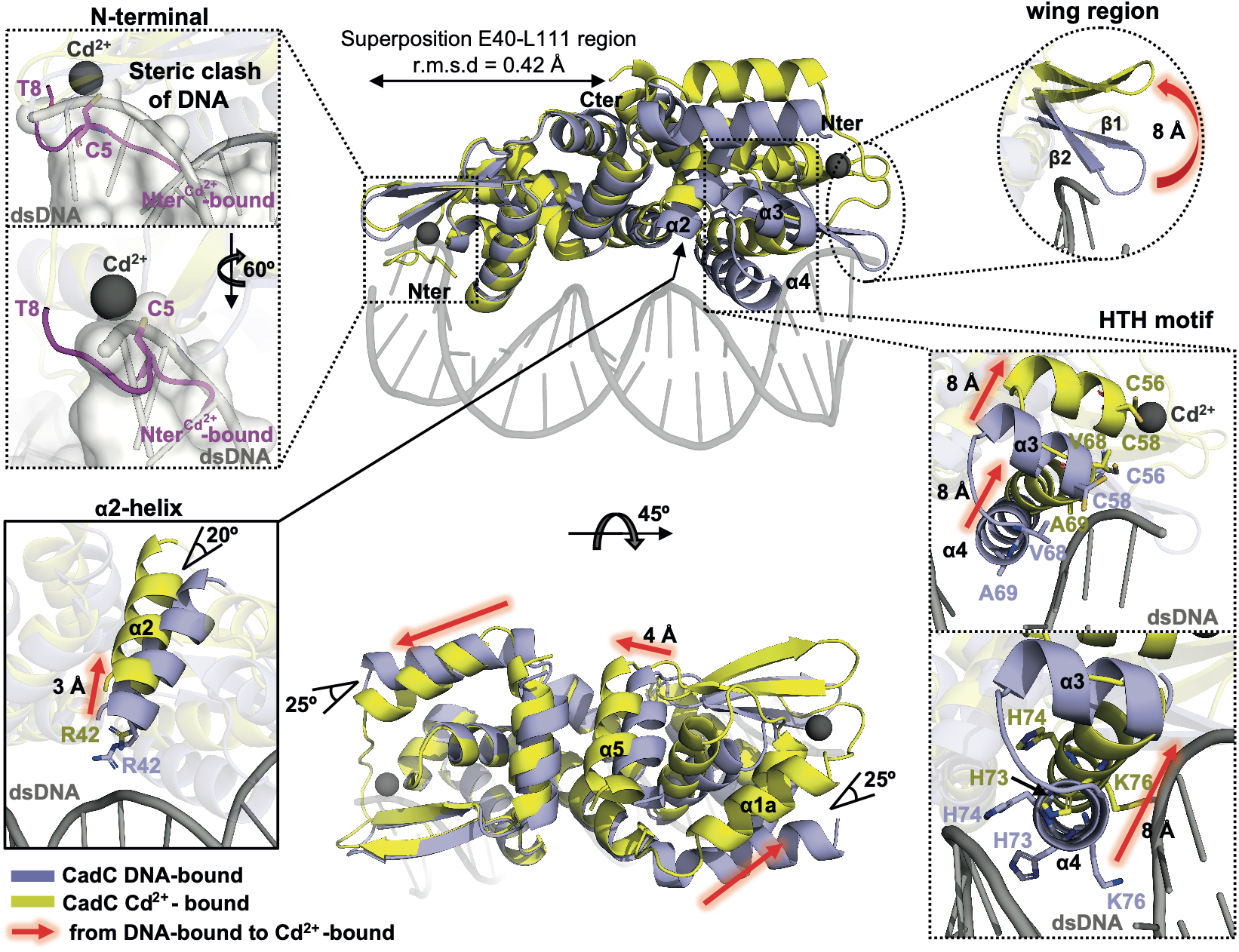
Conformational changes in CadC upon Cd^2+^ binding. Cartoon representation of the superposed structures of DNA-bound CadC (blue) and Cd^2+^-bound CadC (yellow) through residues 40−111 (Cα RMSD: 0.42 Å), illustrating the structural rearrangements associated with Cd^2+^ binding. The N-(Nter) and C-termini (Cter) are indicated. Binding of Cd^2+^ induces pronounced conformational changes, including a repositioning of α-helical elements and shifts within the helix-turn-helix (HTH, bottom left) motif, as well as alterations in the wing region (upper left). Red arrows indicate structural changes between the DNA- and Cd2+-bound CadC structure.

Importantly, the N-terminus (residues M1 to D11), disordered in the apo-CadC structure and truncated to obtain CadC-DNA complex crystals, is ordered in the metal-bound structure, with the model including all residues from D3 onward. Due to its involvement in the coordination of Cd^2+^ through C5, the N-terminus folds back towards the protein and docks in close proximity to helix α4, a structural element implicated in DNA recognition. In structural superposition with the DNA-bound state, the segment M1 to T8 overlaps the position occupied by the DNA duplex, predicting a steric clash (**Figure 6**).

Together, these structural changes are consistent with cadmium binding repositioning key DNA-contacting residues and generating steric incompatibility with the DNA duplex. We therefore propose that Cd^2+^ binding stabilizes a conformation of CadC that is incompatible with productive DNA engagement, thereby promoting dissociation of the CadC-DNA complex.

### Host zinc availability is associated with CadC-dependent infection phenotypes

We previously showed that *cadC* is highly expressed by *Lm* during mouse infection (38) and that deletion of *cadC* results in attenuated virulence (10). To determine whether CadC-dependent virulence is linked to metal sensing under physiologically relevant conditions, we investigated the impact of host metal availability during infection. Although CadC is highly responsive to cadmium *in vitro*, this metal is unlikely to be present at relevant concentrations in host tissues, raising the idea that other metals contribute to CadC regulation during infection. In this context, our data show that zinc binds CadC and modulates CadC-dependent gene expression, supporting the possibility that zinc contributes to CadC regulation in the host environment. We thus assessed the contribution of host zinc availability to CadC function during infection by comparing bacterial burdens in mice fed a control or zinc-restricted diet (**Figure 7A**). Bacterial loads were quantified in the spleen and liver of mice infected with wild-type or Δ*cadC Lm* under control or zinc-deficient dietary conditions. Under a control diet, deletion of *cadC* resulted in a marked and statistically significant reduction of bacterial loads in both spleen and liver as compared with the wild-type strain, confirming the contribution of CadC to virulence (10). In contrast, under the zinc-deficient diet, this difference was no longer detected, as bacterial burdens of the wild-type and Δ*cadC Lm* strains were comparable in both organs (**Figure 7A**). Two-way ANOVA performed separately for each organ revealed a significant diet × genotype interaction in the spleen (F(1,19) = 6.78, p = 0.0175) and liver (F(1,19) = 22.36, p = 0.00015), indicating that the genotype effect depends on dietary zinc status. These findings support host zinc availability as a modifier of the CadC-dependent infection phenotype.

**Figure 7.**
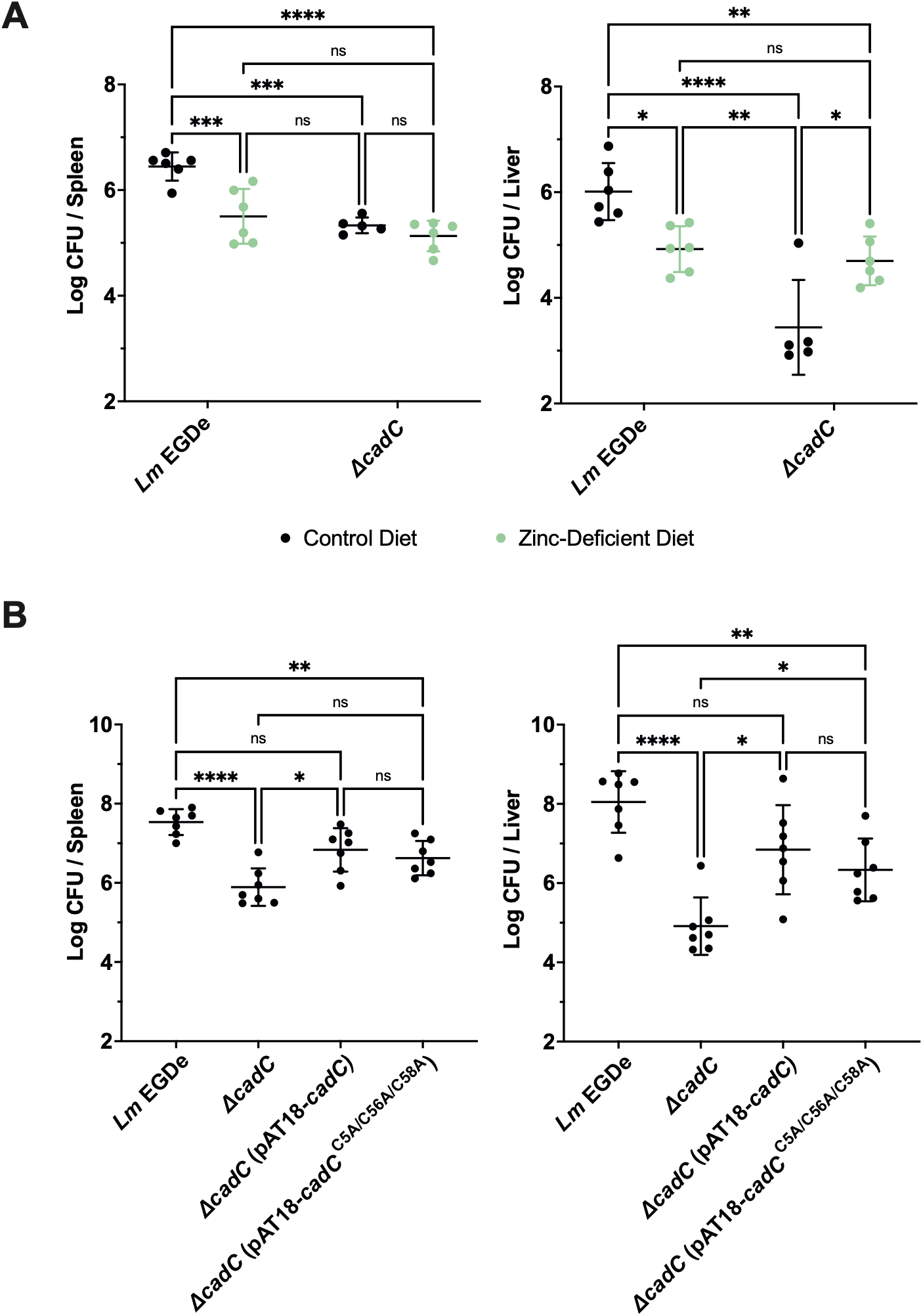
Effect of zinc restriction and CadC cysteine mutations on mouse infection. **(A)** Four-week-old female BALB/c mice (Charles River) were fed either a zinc-deficient or a control diet for 3 weeks prior to intravenous challenge with 10^3^ CFU/animal of *Lm* EGDe or Δ*cadC* (n≥5 per group). **(B)** Seven to eight-week-old female BALB/c mice (Charles River) were intravenously infected with 10^4^ CFUs of *Lm* EGDe, *Lm* Δ*cadC*, *Lm* Δ*cadC* (pAT18-*cadC*) and *Lm* Δ*cadC* (pAT18-*cadC*^C5A/C56A/C58A^). For **(A)** and **(B)**, bacterial burdens in the spleen and liver were determined 72 h post-infection. Data are presented as scatter plots, with each dot representing an individual mouse and horizontal lines indicating mean values. For panel A, log10-transformed CFU/organ values were analyzed separately for spleen and liver by two-way ANOVA with diet and genotype as fixed factors, including the diet × genotype interaction. The interaction was significant in the spleen (p = 0.0175) and liver (p = 0.00015). Wild-type and ΔcadC strains were compared within each diet using Šídák-corrected simple-effects tests. CFU, colony-forming units; ns, not significant, *p* < 0.05 (*); *p* < 0.01 (**); *p* < 0.001 (***); *p* < 0.0001 (****).

To test whether this zinc-dependent effect involved the CadC metal-binding site, we next analyzed the virulence behaviour of a Δ*cadC* strain expressing either wild-type CadC or the CadC cysteine mutant C5A/C56A/C58A, alongside the wild-type or Δ*cadC* strains (**Figure 7B**). The strain expressing the cysteine mutant displayed an intermediate phenotype in both organs: bacterial burdens were significantly higher than those of the Δ*cadC* mutant but remained below wild-type levels. Notably, although the difference did not reach statistical significance, the strain complemented with wild-type *cadC* consistently showed higher bacterial burdens than the strain expressing the cysteine mutant, suggesting a trend towards more efficient functional restoration. Altogether, these results indicate that host zinc availability influences CadC-dependent infection outcomes and are consistent with a contribution of the α3N metal-binding site to CadC-dependent virulence.

## Discussion

How metal binding at a sensory site is converted into release of operator DNA is a central mechanistic question for ArsR/SmtB repressors. The three CadC structures presented here capture the apo, DNA-engaged, and metal-bound endpoints of this regulatory cycle and, together with structure-guided functional analyses, support a DNA-release mechanism in which allosteric remodeling is coupled to incompatibility inferred from structural superposition.

Comparison of the three states suggests a two-component switching mechanism. In the DNA-bound state, the α4 recognition helices, wing regions, and electrostatic contacts are arranged to engage both operator half-sites. Cd²⁺ coordination at the α3N site reorganizes this DNA-binding surface, displacing key contacting residues and shifting the HTH and wing elements away from the duplex. Concurrently, metal coordination orders the N-terminal segment around C5 and positions it across the DNA-binding trajectory. The metal-bound structure therefore supports two complementary barriers to DNA engagement: remodeling of the recognition surface and, in the structural superposition, steric conflict between the ordered N-terminus and the duplex. Together, these changes provide a structural route by which local metal coordination can promote operator release and transcriptional derepression.

CadC adopts the canonical fold of the ArsR/SmtB superfamily, characterized by a wHTH architecture that mediates DNA binding. This overall organization is consistent with other family members (21,22,39–43), supporting a conserved structural framework for DNA recognition, in which positioning of the α4 recognition helix, wing contacts, and electrostatic interactions collectively define operator engagement. In particular, the recognition helix (α4) of CadC inserts into the DNA major groove and establishes base-specific interactions, while additional contacts mediated by the wing region and an extended electrostatic network further stabilize the complex. Importantly, residues outside helix α4 also contribute to DNA binding. We identified R42 and K35/K76 as critical determinants of operator recognition and transcriptional repression.

Our data also highlight the importance of DNA architecture in CadC function. CadC displays a strong dependence on the spacing between palindromic recognition sites, indicating that operator geometry is a key determinant of binding. Both shortening and extension of the canonical spacer length impair binding affinity, indicating that efficient DNA recognition requires precise positioning for simultaneous engagement of both half-sites. When this geometry is disrupted, CadC likely interacts with only a single repeat, resulting in weaker binding. The spacer variants tested here probe the combined effects of half-site distance and helical phasing. Whether the Cd²⁺-bound conformation of CadC could recognize operators with alternative spacing or phasing remains unresolved. This behaviour contrasts with other ArsR/SmtB regulators such as SmtB, which can accommodate more flexible operator arrangements, including binding of multiple dimers or interaction with single half-sites (16,44), suggesting that differences in operator geometry may contribute to regulatory specificity across this family. Thus, variation in operator spacing and symmetry may provide an additional layer of regulatory specificity within the conserved ArsR/SmtB fold.

Analysis of the Cd^2+^-bound structure further reveals an unexpected feature at the α3N metal binding-site. While in *S. aureus* CadC, the α3N site adopts a trigonal S3 coordination geometry with Pb^2+^ (C7, C58 and C60) and rearranges to a tetrahedral S4 geometry upon Cd^2+^ or Bi^3+^ binding through recruitment of an additional cysteine (C11) (29,45), *Lm* CadC instead coordinates cadmium using three cysteines (C5, C56 and C58) and one methionine (M92). Notably, CadC retains the conserved C9 residue that typically participates in metal coordination in α3N-type sites (29,40,46), yet in the structure this residue does not contribute to Cd^2+^ binding. There is limited structural information available for ArsR/SmtB regulators with metals bound at α3N-type sites. Current examples are restricted to ArsR homologues from *Acidithiobacillus ferrooxidans* and *Corynebacterium glutamicum*, where As^3+^ is coordinated in S3 configuration (13). Our structure therefore reveals unexpected plasticity at an α3N-type site: despite retention of the conserved C9, a methionine replaces the canonical cysteine ligand in the observed Cd²⁺ coordination sphere. This finding expands the known coordination chemistry of ArsR/SmtB metal sensors and shows that sequence conservation alone does not necessarily predict the realized ligand set. Given the limited number of structurally characterized α3N-type sites, this finding suggests that metal coordination within ArsR/SmtB regulators is more flexible than previously appreciated (5). Whether this combination of conformational remodeling and steric control is shared by other ArsR/SmtB metalloregulators remains to be established.

At the mechanistic level, CadC displays metal-specific sensing behavior. Cadmium binds apo-CadC with higher apparent affinity than zinc, indicating that CadC is preferentially tuned toward cadmium recognition. This selectivity is consistent with the broader principles of metal discrimination in metalloregulatory proteins, where metal preference is not dictated exclusively by the first coordination sphere, but also by coordination geometry, second-shell interactions, protein conformational dynamics, and the capacity of a given metal ion to stabilize the regulatory allosteric state (47,48). For Cd²⁺, within the adopted two-event model, the large separation between the fitted macroscopic binding constants is consistent with apparent negative cooperativity, although alternative models cannot be formally excluded. Zn²⁺ also displays two fitted binding events, but their similar apparent affinities preclude a strong assignment of cooperativity. A similar mechanism has been described for *S. aureus* CadC (47), suggesting that this behavior may be conserved within this protein subgroup. By contrast, in CzrA, zinc binding occurs at the distinct α5 site and follows a thermodynamically nonequivalent two-site mechanism (49), highlighting that the structural context of the metal-binding site defines the mechanistic basis of allostery in ArsR/SmtB regulators. The apparent negative cooperativity may also involve changes in conformational dynamics. Dynamics-driven negative cooperativity has been described for the cyclic AMP receptor protein (CAP), where ligand binding alters conformational fluctuations and the entropic cost of subsequent binding events (50,51). Whether a related mechanism contributes to sequential metal binding by CadC remains to be determined.

Beyond metal homeostasis, several ArsR/SmtB regulators have been implicated in virulence-associated functions, underscoring the adaptability of this protein family in integrating environmental cues with pathogenic processes. Examples include *Brucella abortus* ArsR2, which regulates the T4SS activator VjbR and is required for full virulence in mice (52), *Brucella suis* S2 ArsR6, which controls the stress-associated outer membrane protein Omp25D (53), and the ArsR-like regulator Rv2034 of *Mycobacterium tuberculosis*, which activates the hypoxia-responsive DosR regulon (54). However, the physiological signals that activate such regulators during infection have remained largely unclear.

Although CadC is most responsive to cadmium *in vitro*, this metal is unlikely to be encountered at physiologically relevant levels in mammalian tissues (55). Our data show that host zinc availability modifies CadC-dependent infection outcomes. Despite its lower affinity for CadC compared to cadmium, zinc exhibits measurable binding, induces transcription from CadC-regulated promoters in bacterial cells, and under zinc limitation the difference between the WT and Δ*cadC* strains was no longer detected. In addition, mutation of the α3N cysteine residues resulted in partial loss of CadC function *in vivo*, consistent with a contribution of this metal-binding site to CadC activity under physiological conditions. Together with the dietary zinc experiments, these findings establish an association between host zinc availability and CadC-dependent infection phenotypes, although direct zinc sensing by CadC during infection remains to be established. Notably, this relationship differs from that described in several other pathogens, including *Trypanosoma cruzi, Heligmosomoides polygyrus, Streptococcus pneumoniae, Acinetobacter baumannii* and *enteroaggregative Escherichia coli*, where dietary zinc restriction enhances infection or virulence (56–60). This dual role, coupling metal sensing with regulation of virulence-associated genes, distinguishes CadC from many ArsR/SmtB homologs that remain restricted to metal homeostasis, and highlights how this family can be adapted to integrate environmental sensing with broader regulatory networks (30,61).

Together, the apo, DNA-bound, and Cd2+-bound structures define how a metal-responsive ArsR/SmtB repressor traverses between transcriptionally repressive and derepressive states. CadC recognizes its operator through coordinated major-groove, wing, and backbone interactions, whereas metal coordination stabilizes a DNA-incompatible state through coupled remodeling of these recognition elements together with predicted steric incompatibility of the ordered N-terminus with the DNA duplex. The observed noncanonical Cys3-Met coordination geometry further expands the chemical repertoire of α3N metal-sensing sites. In *Listeria*, this regulatory module operates beyond cadmium detoxification, and host zinc availability is associated with changes in CadC-dependent infection phenotypes. More broadly, the work provides a structural mechanism for converting metal sensing into DNA release in ArsR/SmtB transcriptional regulation.

## Methods

### Bacterial strains and growth conditions

Strains and plasmids used in this study are listed in **Supplementary Table S1**. *Lm* EGDe and *E. coli* strains were routinely cultured in Brain Heart Infusion (BHI, Difco) and/or Chemical Defined Medium (MM) (62), and Lysogenic Broth (LB), respectively, at 37 °C aerobically with shaking. BHI and LB agar (Difco) plates were used for growth on solid media. When appropriate, media were supplemented with antibiotics at the following concentrations: ampicillin 100 μg/ml, chloramphenicol (Cm) 20 μg/ml, erythromycin (Ery) 150 μg/ml, kanamycin (Kan) 50 μg/ml and tetracycline 12.5 μg/ml for *E. coli* strains; and Cm 7 μg/ml and Ery 5 μg/ml for *Lm* strains.

### Cloning

The native *Lm EGDe cadC* sequence was fused to a 6xHis-tag and cloned into pET28b, generating pET28b(*cadC*-6His) (10). The truncated variant of CadC comprising the residues D11-L111 (CadC^D11-L111^) was generated by high-fidelity PCR amplification (ACCUZYME Mix, Bioline) using *Lm* EGDe genomic DNA as template, with oligonucleotides DM83 and DM84 listed in **Supplementary Table S2**. The amplified fragments were subsequently cloned into the pET28b in frame with an C-terminal 6xHis-tag.

### Site-directed mutagenesis

CadC amino acid substitutions (CadC^C56A/C58A^, CadC^C5A/C56A/C58A^, CadC^K35A^, CadC^R42A^, CadC^V68G^, CadC^A69L^, CadC^H73A/H74A^, CadC^K76A^ and CadC^K35A/K76A^) were generated by site-directed mutagenesis, using pET28b(*cadC*-6His) as template and the oligonucleotides listed in **Supplementary Table S2**. Parental methylated DNA was digested with *Dpn*I (Thermo Fisher Scientific).

### Expression and purification of recombinant proteins

Full-length His-tagged native CadC, along with its mutant variants were expressed in *E. coli* BL21(DE3). Protein expression was induced at OD_600nm_∼0.6 at 37 °C with the addition of 0.5 mM IPTG for 3 h. Bacteria were harvested by centrifugation at 4000 x g for 30 min, resuspended in lysis buffer (50 mM MOPS pH=7.5, 500 mM NaCl, 1 mM TCEP and 10 mM Imidazole) and lysed by sonication, followed by centrifugation at 36000 x g for 45 min. Recombinant proteins were captured from the supernatant by nickel-affinity chromatography (Ni-NTA agarose, ABT), and eluted with 50 mM MOPS pH 7.5, 500 mM NaCl, 1 mM TCEP, 100-400 mM Imidazole. Imidazole was removed by dialysis (Spectra/Por Dialysis Membrane MWCO: 6-8 kD, SpectrumLabs) in a buffer containing 50 mM MOPS pH 7.5, 200 mM NaCl and 1 mM TCEP. Eluted proteins were concentrated using 10 kDa molecular-weight cut-off centrifugal concentrators (Vivaspin, Sartorius Stedim Biotech) and further purified by size-exclusion chromatography (Superdex200 Increase 10/300 GL, Cytiva). Recombinant proteins were analyzed by SDS-PAGE and purities determined by densitometry of Coomassie Blue-stained gels (Image Lab Software, BioRad). Obtained recombinant proteins were ≥90% pure (**Supplementary Figure S2**). Protein concentrations were determined using Bradford assay (Bio-Rad). For infrared quantification, protein samples were applied to Direct Detect Assay-free membrane cards and analyzed by Fourier-transform infrared spectroscopy based on the absorption of peptide-bond amide groups using Direct Detect Infrared Spectrometer (Merck/Sigma-Aldrich). Molecular weights were calculated with the ExPASy ProtParam tool. Purified proteins were flash-frozen in liquid nitrogen and stored at -80 °C.

### Transcriptional fusion reporters

The pAD-P*actA*-GFP vector (63), was used as template for PCR amplification (ACCUZYME Mix, Bioline) of a fragment containing the *actA* promoter fused to *gfp*-encoding sequence, using the oligonucleotides pPL2_SacI_Fw and pPL2_SacI_Rv (**Supplementary Table S2**). The PCR product was digested with *SacI*-HF (NEB) and re-ligated into the *Sac*I-digested and dephosphorylated (FastAP thermosensitive alkaline phosphatase, Thermo Fisher Scientific) integrative plasmid pPL2-mCherry (64). To generate *cadC* and *cadA* transcriptional reporters, the resulting plasmid (pPl2-mCherry-P*actA*-GFP) was digested with *Sma*I-HF and *BamH*I-HF (NEB), allowing the excision of the *actA* promoter sequence. *cadC* and *cadA* promoter sequences were PCR-amplified from *Lm* EGDe genomic DNA using oligonucleotides P*cadC*_SmaI_Fw and P*cadC*_BamHI_Rv, or P*cadA*_SmaI_Fw and P*cadA*_BamHI_Rv, respectively. The PCR fragments were digested with *Sma*I-HF and *BamH*I-HF (NEB) and introduced into the pPl2-mCherry-P*actA*-GFP digested with the same restriction enzymes. The generated constructs were confirmed by DNA sequencing. Plasmids were electroporated in *Lm* EGDe wild-type and Δ*cadC*, and transformants selected at 37°C in BHI agar-Cm7. Integration into the *Lm* chromosome was confirmed by PCR using the oligonucleotides PL95 and PL102.

### *Lm* strains carrying *cadC* point mutations

To generate the *Lm* strains expressing specific *cadC* mutant variants, mutations were introduced in the non-integrative pAT18 vector (65). The *cadC* promoter sequence was amplified by PCR using P*cadC* Fw and DM66 oligonucleotides. The *cadC* coding sequences were amplified by PCR: the wild-type *cadC* from genomic DNA, and the mutated *cadC* versions from pET28b-based constructs, carrying either C56A+C58A (*cadC*^C56A/C58A^), C5A+C56A+C58A (*cadC*^C5A/C56A/C58A^), R42A (*cadC*^R42A^), K35A+K76A (*cadC*^K35A/K76A^) and H73A+H74A (*cadC*^H73A/H74A^) substitutions, using oligonucleotides DM67 and DM68. The PCR products of the *cadC* promoter and the different *cadC* sequences were fused by Splicing-by-Overlap-Extension (SOE) PCR, using the oligonucleotides P*cadC* and DM68. The P*cadC*-*cadC* fragments were cloned at the *Sma*I and *BamH*I restriction sites of a digested and dephosphorylated pAT18. The constructs were verified by DNA sequencing. Generated pAT18 harboring mutated *cadC* variants were electroporated to introduce into *Lm* Δ*cadC*::mCherry-P*cadA*-GFP (*Lm* Δ*cadC* P*cadA*).

### Fluorescence Anisotropy

DNA-binding studies were performed using a 1:1 (molar:molar) mixture of fluorescein-labeled complementary oligonucleotides containing the CadC-box (**Supplementary Table S2**). Oligonucleotides annealing was achieved by heating the mixture at 95°C for 5 min and then cooling at 1°C/min until 25°C. Anisotropy measurements were performed in a plate-format in a filtered-buffer containing 50 mM MOPS pH 7.0, 300 mM NaCl and 1 mM TCEP with 5’-fluorescein-labeled DNA (75 nM) and titrated with increasing concentrations of CadC, or with the DNA-CadC complex mixture (1 µM of 5’-fluorescein-labelled DNA and 4 µM of CadC) titrated with increasing concentrations of CdCl_2_. Titration mixtures were prepared in 96-flat bottom nontreated black well plates (Thermo Fisher Scientific) and incubated for 2 h in the dark at room temperature. Fluorescence anisotropy was read on a plate reader (BioTek Synergy 2) using excitation and emission filters 485/20 and 528/20 nm, respectively. Experimental data were fitted in OriginPro8.5 using a Hill equation 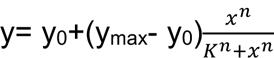 where *y* is the measured anisotropy, *y_0_* is the baseline anisotropy, *y_max_* is the anisotropy at saturation, *x* is the protein concentration, *K* is the apparent half-saturation concentration (K1_/2_) derived from the empirical Hill fit, and n is the Hill coefficient. Experimental data from metal cation titrations of DNA-CadC complexes were fitted in DynaFit, using a competition binding model: DNA-Protein ⇌ DNA+Protein (Kd), Protein+Cd^2+^ ⇌ Protein-Cd^2+^ (Kdd) where the constant of dissociation for the DNA-Protein complex was fixed (values shown in **Supplementary Tables S3 and S4**) and the constant of dissociation of the Protein-Cd^2+^ varied. In graphical representations using a logarithmic scale, the data point corresponding to 0 protein concentration was plotted as 0.001 or 0.00001 (in the same concentration units); this value was included in curve fitting.

### Bacterial growth and gene transcription analysis

*Lm* strains at OD_600nm_ ∼0.8 were diluted 1:20 in MM in black 96-well microplates with clear flat-bottom (Greiner) at a final volume of 200 μl/well. Plates were incubated at 37 °C with shaking. After 5 h, increasing concentrations of CdCl_2_ or ZnCl_2_ (Sigma-Aldrich) were added and cultures re-incubated in the same conditions. Growth (OD_600nm_) and fluorescence were monitored every 20 min using a CLARIOstar Plus plate reader (BMG Labtech). Fluorescence was measured under the following parameters: mCherry (Excitation 570/15 nm, Emission 620/20 nm; gain = 1500/2000) and GFP (Excitation 470/15 nm, Emission 515/20 nm; gain = 1000/1500). GFP fluorescence in each well was normalized by subtracting the GFP background signal of the control strain (*Lm*::mCherry), and by dividing by the mCherry signal from the same well.

### Size Exclusion Chromatography (SEC)

SEC was performed using a High-Resolution Liquid Chromatography System (ÄKTA Purifier 10, Cytiva) equipped with a Superdex 200 Increase 10/300 GL column (Cytiva). The column was equilibrated with EQ Buffer (50 mM MOPS pH 7.5, 100 mM NaCl, and 1 mM TCEP), and operated at a flow rate of 0.5 ml/min. CadC (50 μM), FAM-labeled DNA (25 μM), and CadC-DNA complex with 25 μM CdCl_2_ were prepared in EQ buffer, incubated at room temperature for 2 h, and injected in 200 μl sample loops. Elution profiles were monitored by UV absorbance at 280 nm (protein) and 260 nm (DNA). Fractions were collected throughout the run and Fluorescein (FAM) fluorescence measured using a CLARIOstar Plus plate reader (BMG Labtech) (excitation: 483/14 nm, emission: 530/30 nm). Fractions were also analyzed by SDS-PAGE (17 % acrylamide).

### Crystallization and structure determination

Initial crystallization conditions for CadC, CadC-Cd^2+^ and CadC-DNA complex were determined at 20 °C using commercial sparse-matrix crystallization screens. Sitting-drop vapor diffusion experiments were set up in MRC 96-Well 2-Drop Crystallization plate (Molecular Dimensions) using an Oryx 4 crystallization robot (Douglas Instruments). Apo-CadC crystals were obtained by mixing equal volumes (0.15 μl) of CadC protein solution (at 10 mg/ml in 50 mM MOPS pH 7.5, 200 mM NaCl, 1 mM TCEP) and crystallization solution (8 % v/v Tacsimate pH 5.0 and 20 % w/v PEG 3350), equilibrated against 70 μl reservoir solution. For data collection, crystals were cryo-protected by sequential transfer into their crystallization condition with increasing concentrations of glycerol (up to 30 % v/v) and then flash-frozen in liquid nitrogen (66). The CadC structure with bound Cd^2+^ was obtained from a crystal grown with equal volumes (0.15 µl) of CadC protein solution (at 10 mg/ml in 50 mM MOPS pH 7.5, 200 mM NaCl, 1 mM TCEP and with 1 mM CdCl_2_) and crystallization solution (100 mM HEPES pH 7.5, 10 % v/v 2-propanol, 20 % w/v PEG 4K, 15 % v/v glycerol). The crystal was directly frozen in liquid nitrogen. Prior to crystallization assays with DNA, 1 mM of CadC^D11-L111^ was incubated 2 h at room temperature with 250 µM of the ATTCAAACAAACACTTGAAT double strand DNA box (Thermo Fisher Scientific). The DNA duplex length was selected based on AlphaFold 3 (31) models of the CadC-DNA complex to define the minimal DNA region required for binding while minimizing unbound terminal DNA segments that could introduce conformational flexibility and hinder crystal formation. Then, 0.15 µl of this solution was mixed with an equal volume of crystallization condition composed of 0.5 mM of each Oxometalate (sodium chromate tetrahydrate, sodium molybdate dihydrate, sodium tungstate dihydrate and sodium orthovanadate), 100 mM BES/TEA pH 7.5, 15 % w/v PEG 3K, 20 % v/v 1,2,4-butanetriol, 1 % w/v NDSB 256. Crystals were directly frozen in liquid nitrogen.

Datasets were collected on ID30b (CadC and CadC-Cd structure) and MASSIF-3 (DNA-bound structure) beamlines (67,68) at ESRF Synchrotron (Grenoble, France). The CadC apo-structure was determined at 3.0 Å resolution by molecular replacement using PHASER (69) with a CadC model predicted by AlphaFold 3 (31) as a search template. Several cycles of model building and refinement were performed using COOT (70) and Phenix.refine (71–73). Refinement was performed using standard stereochemical restraints. Given the presence of six CadC molecules in the asymmetric unit, non-crystallographic symmetry (NCS) restraints were applied between equivalent chains during refinement. Structures of CadC in complex with Cd^2+^ and DNA were subsequently solved by molecular replacement using the refined apo-structure as a search model. The CadC-Cd and CadC-DNA complexes were refined to resolutions of 2.0 Å and 1.7 Å, respectively, following multiple rounds of model building and refinement.

For the CadC-DNA complex, data processing was performed in the symmetry space group C2, in which the asymmetric unit comprises a CadC subunit bound to a single DNA strand, with the other dimer subunit and complementary DNA strand generated by crystallographic symmetry. Data collection and refinement statistics are indicated in **Table 1**.

### Animal infections

Animals were acclimatized for 7 days prior to dietary intervention and infection. Four-week-old BALB/c mice (Charles River Laboratories) were fed for three weeks either a zinc-restricted diet (E15571-24, Ssniff) or a control diet (E15712-04, Ssniff). Blood samples were collected from the posterior vena cava and Zn^2+^ quantified by inductively coupled plasma mass spectrometry (ICP-MS). Measured zinc levels were lower in mice fed a zinc-restricted diet (793 ± 65 μg/l) compared to those fed a control diet (1091 ± 193 μg/l). Mice (n≥5 per group) were intravenously infected via tail vein injection with 10^3^ colony-forming units (CFUs) of *Lm* EGDe wild-type or Δ*cadC* mutant strain as previously described (74).

Additionally, 7-8-week-old female BALB/c mice (Charles River Laboratories) were intravenously infected by inoculation of 10^4^ CFUs of *Lm* EGDe, *Lm* Δ*cadC*, *Lm* Δ*cadC* (pAT18-*cadC*) and *Lm* Δ*cadC* (pAT18-*cadC*^C5A/C56A/C58A^). At 72 h post-infection, mice were euthanized by CO_2_ inhalation followed by cervical dislocation. Organs were collected and homogenized in PBS and homogenates were serially diluted and plated on BHI agar plates. The day after, CFUs were quantified. Sample sizes were selected based on prior studies using comparable *Listeria infection* models and animals were randomly assigned to groups. Welfare was assessed daily by measuring body weight and inspecting appearance (piloerection, posture), behaviour, hydration, and respiration movement. Mice were housed at the i3S animal facility under specific pathogen-free conditions in high-efficiency particulate air (HEPA)-filtered cages, maintained on a 12 h light/dark cycle, and provided with sterile chow and autoclaved water ad libitum. Animal procedures followed the guidelines of the European Commission for the handling of laboratory animals (Directive 2010/63/EU) and the Portuguese legislation for the use of animals for scientific purposes (Decreto-Lei 113/2013). All procedures were approved by the i3S Animal Ethics Committee and Direcção Geral de Alimentação e Veterinária (license 006564).

### Isothermal Titration Calorimetry (ITC)

Wild-type (WT) and mutant CadC recombinant proteins were dialysed overnight against 50 mM HEPES pH 7.5, 200 mM NaCl and 1 mM TCEP and ligands were dissolved in the dialysis buffer. Both proteins and ligands were filtered and degassed. ITC titrations were carried out on a MicroCal VP-ITC calorimeter, at 15 °C, using 200 µM of divalent metal cation (syringe) and 10 µM of CadC (reaction cell). The initial injection of metal cation solution was 2 µl, followed by 14 injections of 4 µl and 34 additional injections of 10 µl, in 500 s intervals. Heat integrations were performed using NITPIC, and data were fitted using SEDPHAT. The resulting SEDPHAT output files were then directly imported into Gussi software for the plot of ITC data (75). Prior to analysis, the corresponding blank titration (200 µM CdCl_2_ into 50 mM HEPES pH 7.5, 200 mM NaCl, 1 mM TCEP) was subtracted from the raw data. For CadC, a two-site binding model (A+B+B ⇌ (AB)+B ⇌ ABB) with two symmetric binding sites and macroscopic equilibrium constants was used. For CadC^C5A/C56A/C58A^, data were fitted using a simple 1:1 binding model (A+B ⇌ AB). Reported dissociation constants (Kd) were calculated with 68.3 % confidence intervals (68.3 % Cl).

### Thermal Shift Assay

Protein stability was assessed by thermal shift assays, in triplicate. Reactions contained 5 μM of WT or mutant CadC proteins in 50 mM MOPS pH 7.5, 200 mM NaCl, 1 mM TCEP, and 5x SYPRO Orange (Supelco) in a final volume of 50 μL. Samples were equilibrated at 25 °C for 31 s, followed by 140 heating cycles (30 s hold with a temperature increase of 0.5 °C per cycle). Fluorescence corresponding to the SYPRO Orange signal was measured in CFX96 PCR System (Bio-Rad) over time, using the HEX channel (excitation 530 nm/ emission 555 nm), and melting temperatures (Tm) were determined using CFX Maestro software (Bio-Rad) (76).

### Cytosolic *Lm* protein extracts and Western Blot analysis

CadC expression in the different *Lm* CadC mutant strains was assessed in cytosolic protein extracts. Briefly, 100 ml cultures were grown to exponential-phase (OD_600_ of ∼0.5), supplemented with 384 μM of CdCl_2_, and further incubated at 37 °C until reaching an OD_600_ of ∼1. Bacteria were harvested by centrifugation (3200 xg, 15 min, 4 °C), resuspended in 500 µl of lysis buffer (20 mM Tris-HCl pH 7.4 supplemented with Protease Inhibitor Cocktail (cOmplete™, Roche), and mechanically disrupted by bead beating in a FastPrep apparatus (5 cycles of 60 s at 6.5 m/s). Lysates were clarified by centrifugation (850 xg, 10 min) and soluble proteins were obtained from the supernatants after further centrifugation (16000 xg, 15 min). Protein extracts were resolved by SDS-PAGE on a 15% polyacrylamide gel and transferred onto nitrocellulose membrane for 10 min (25 V constant, 1.3 A) using a Trans-Blot Turbo Transfer System (Bio-Rad). Membranes were stained with Ponceau S before blocking in 5 % w/v bovine serum albumin (BSA, Sigma Aldrich) for 1h at room temperature in Buffer A (200 mM Tris-HCl pH = 7.4, 0.9% (w/v) NaCl, 0.1% (v/v) Tween-20). Membranes were incubated overnight at 4°C with the primary antibody rabbit polyclonal anti-CadC (1:500; (30)) diluted in Buffer A with 2.5% (w/v) BSA. After three washes in Buffer A, membranes were incubated for 1 h at room temperature with anti-rabbit horseradish peroxidase (HRP)-conjugated secondary antibody (1:2000; P.A.R.I.S Biotech), in Buffer A with 2.5% (w/v) BSA. Following three additional washes, signals were detected using the Pierce® ECL Western Blotting Substrate (Thermo Fisher Scientific) and digitally acquired in a ChemiDoc XRS+ system (Bio-Rad Laboratories).

### Statistical analyses

Statistical analyses were conducted using Prism 10 (Graphpad Software). For the dietary zinc experiment, log10-transformed CFU/organ values were analyzed separately for spleen and liver using two-way ANOVA with diet and bacterial genotype as fixed factors, including the diet × genotype interaction. Following the interaction analysis, prespecified simple-effects comparisons between wild-type and Δ*cadC* strains within each diet were performed using Šídák’s correction for multiple comparisons. To evaluate infectious capacity among strains (*Lm* EGDe, *Lm* Δ*cadC*, *Lm* (pAT18-*cadC*) and *Lm* Δ*cadC* (pAT18-*cadC* C5A/C56A/C58A)), data normality was evaluated using the Shapiro-Wilk test, and homogeneity of variances was assessed using the Brown-Forsythe test. A one-way ANOVA was applied to compare differences among bacterial strains. Post hoc multiple comparisons were carried out using Dunnett’s T3 test to evaluate pairwise differences between groups. Data are presented as individual values with mean ± standard deviation (SD). Differences with a calculated p-value >0.05 were considered non-significant and statistically significant differences were noted as follows: *p < 0.05; **p < 0.01; ***p < 0.001; ****p < 0.0001.

## Supplementary data

Supplementary data File

## Acknowledgements

We thank Liliana Oliveira (Cell Activation & Gene Expression, i3S) for assistance with blood sample collection for zinc quantification. We acknowledge the European Synchrotron Radiation Facility (ESRF) for provision of beamtime and thank the staff for their support during data collection. The authors also acknowledge the support of the Animal Facility, Biochemical and Biophysical Technologies, X-ray Crystallography and BioSciences Screening i3S scientific platforms, member of the PT-OPENSCREEN (NORTE-01-0145-FEDER-085468) and PPBI (PPBI-POCI-01-0145-FEDER-022122) (https://www.i3s.up.pt/scientific-platforms.php).

## Author Contributions

D.M., J.L., J.M.-C. and D.C. conceived the study. D.M., J.L., R.P., S.S., J.M.-C. and D.C. conceived and designed the experiments. D.M., A.A. and J.L. performed the experiments and data acquisition. D.M., J.L., R.P., S.S., J.M.-C. and D.C. analyzed and interpreted the data. J.M.-C. and D.C. had management as well as coordination responsibility for the execution of the research work and contributed to the acquisition of the financial support and resources. D.M. and J.L. prepared the figures. D.M., J.L., J.M.-C and D.C wrote the manuscript with contribution by R.P. and S.S. All authors reviewed critically the manuscript and approved the final version of the manuscript.

## Data availability

Coordinates were deposited at the PDB under accession codes 9T6R (apo-CadC), 31RT (CadC-Cd^2+^), and 31NU (CadC-DNA). The corresponding diffraction images have been deposited at the SBGrid Data Bank:

- apo-CadC, https://doi.org/10.15785/SBGRID/1259;
- CadC-Cd^2+^, https://doi.org/10.15785/SBGRID/1260;
- CadC-DNA, https://doi.org/10.15785/SBGRID/1261.

Raw data underlying this article were deposited at https://data.4tu.nl/private_datasets/Qz4hMpqmFCvRvmG1RzVXez1dcxZpGScVLMZY4Q67BFM.

## Supplementary Information

**Supplementary Figure S1.**
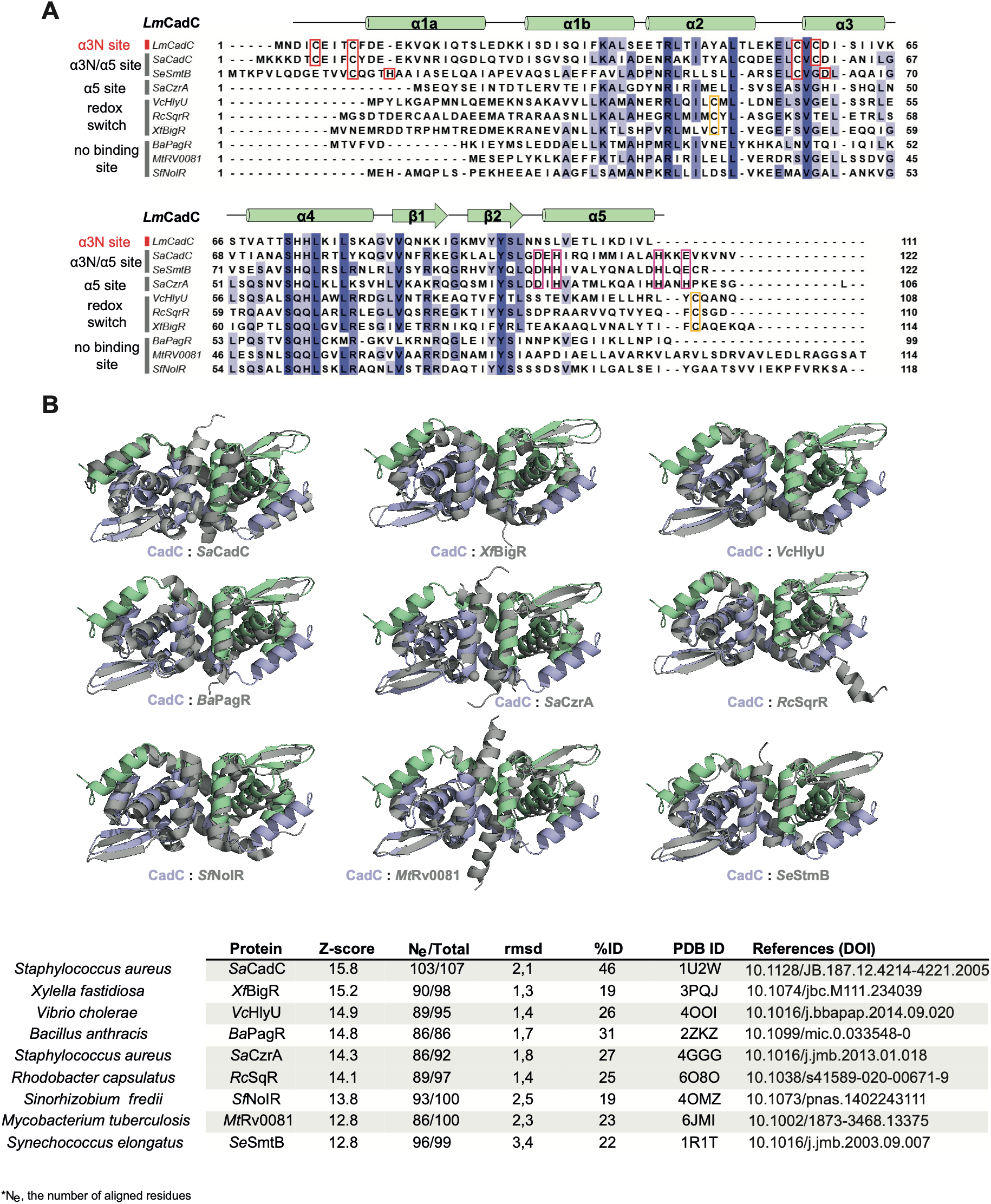
Sequence and structure comparison of CadC and ArsR/SmtB family transcription factors. **A)** Multiple sequence alignment of *Listeria monocytogenes* CadC (Q8Y817), *Staphylococcus aureus* cadmium resistance transcriptional regulator CadC (P20047)*, Synechococcus elongatus* transcriptional repressor SmtB (P30340), *Staphylococcus aureus* CzrA (Q2FWB0)*, Vibrio cholerae* transcriptional activator HlyU (P52695)*, Rhodobacter capsulatus* transcriptional regulator SqrR (D5AT91)*, Xylella fastidiosa* biofilm growth-associated repressor BigR (Q9PFB1)*, Bacillus anthracis* transcriptional repressor PagR (O31178)*, Mycobacterium tuberculosis* Rv0081 (P9WMI7) *and Sinorhizobium fredii* NolR (Q83TD2). α-Helices and β-strands of CadC are indicated above the alignment. **B)** Pairwise structural comparisons of CadC crystal structure (coloured as in Figure 1B) and its closest structural homologues. Proteins structurally related to CadC were identified using the DALI server (http://ekhidna.biocenter.helsinki.fi/dali_server/). Results are summarized in the bottom table. RMSD, root mean square deviation; Ne, the number of aligned residues; %ID, percentage of sequence identity.

**Supplementary Figure S2.**
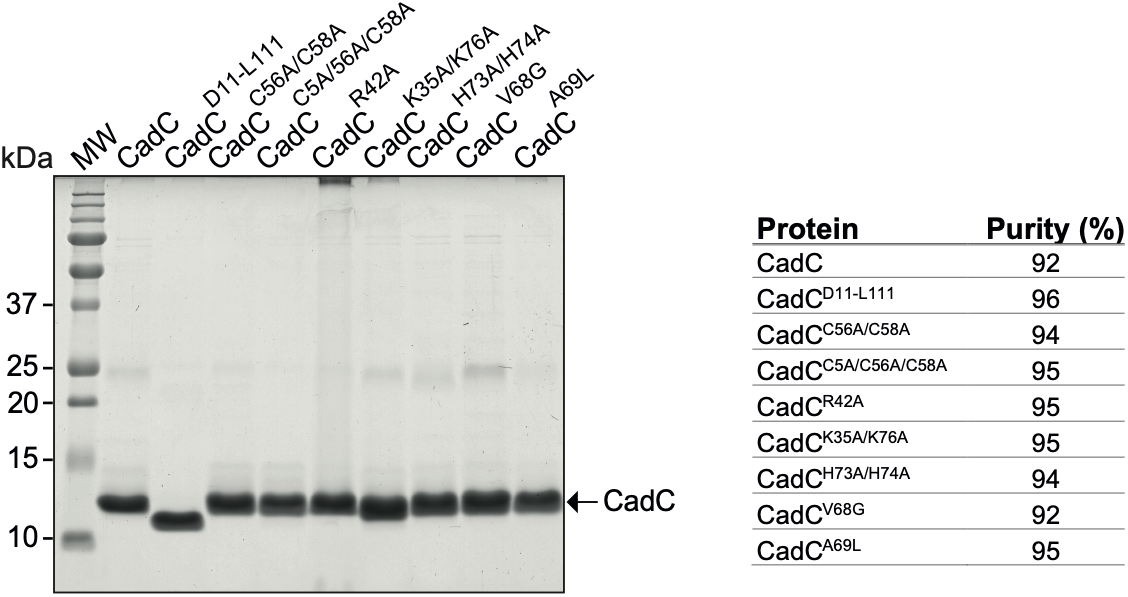
SDS–PAGE analysis of purified recombinant proteins. Five micrograms of each recombinant protein were loaded on a 15% SDS–PAGE gel and visualized by Coomassie Brilliant Blue staining. Lane M, molecular mass markers (kDa; sizes indicated on the left). Purified recombinant proteins are identified above the lanes. The purity of each sample was estimated by densitometry using ImageJ and is summarized in the table.

**Supplementary Figure S3.**
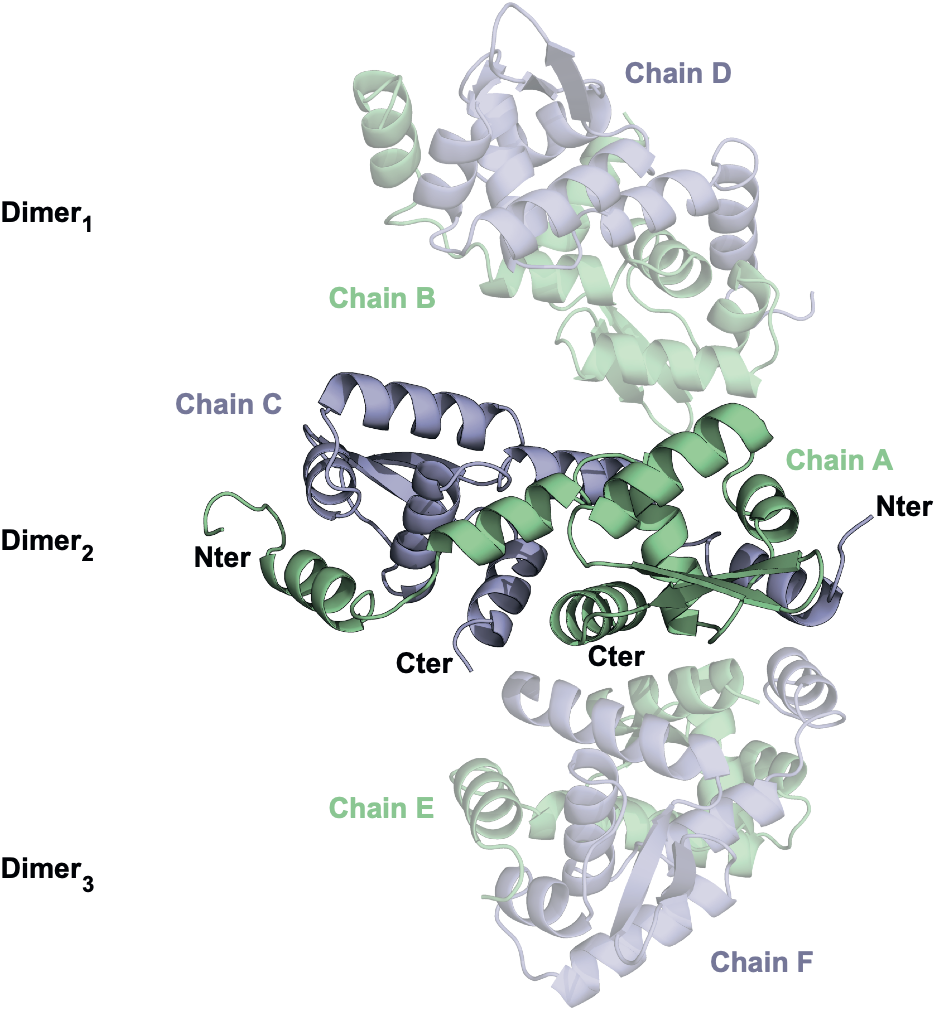
Asymmetric unit of CadC. Cartoon representation of the CadC crystal packing. The asymmetric unit is composed of six molecules of CadC forming three homodimers (Dimer1, Dimer2 and Dimer3).

**Supplementary Figure S4.**
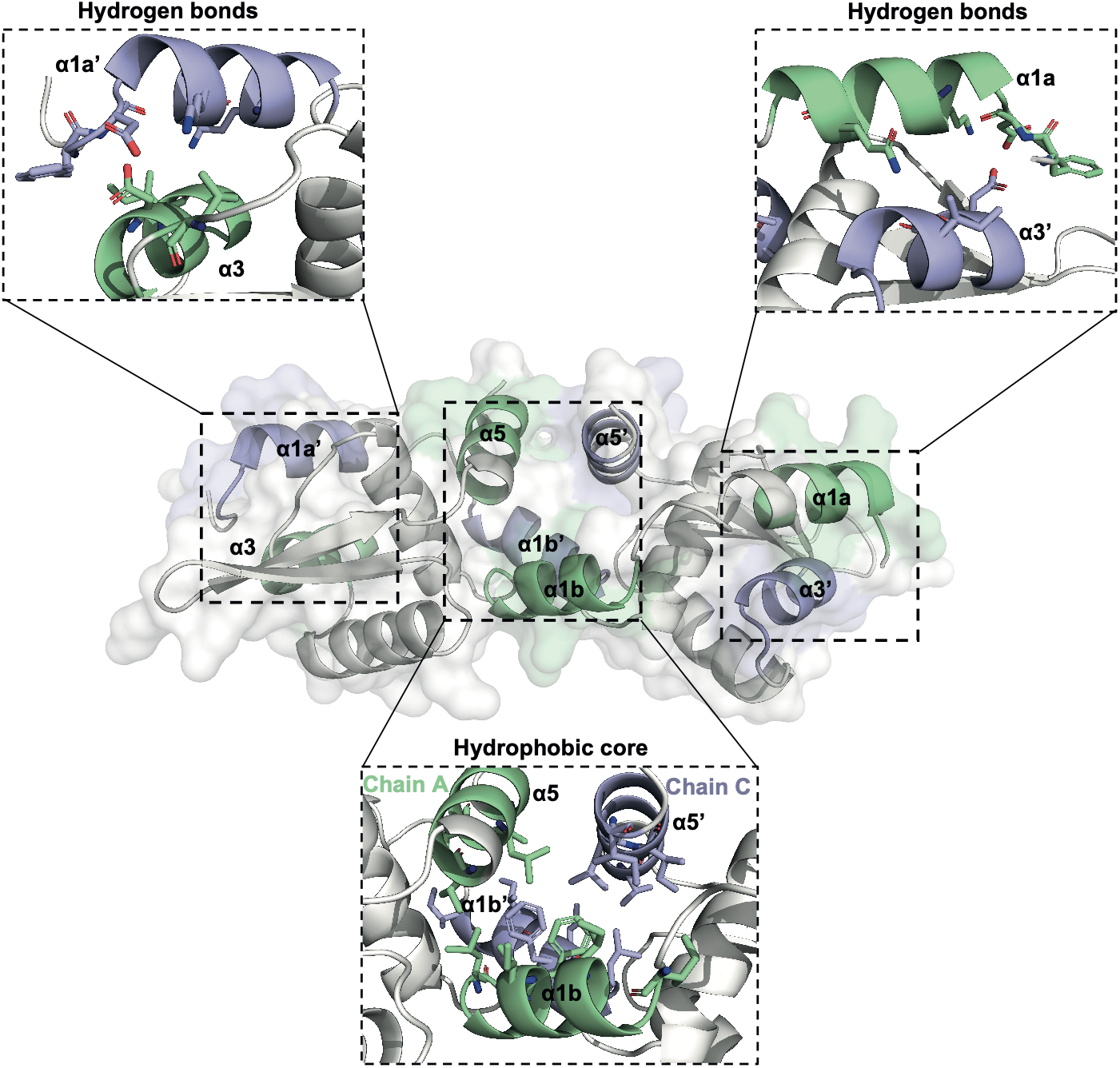
CadC dimer. The dimer interface of CadC is formed by coiled-coil interactions between the α1a and α3 helices (top insets). The dimer is stabilized by a central hydrophobic core involving α1b and α5 helices (bottom inset). The interacting residues were identified by the PDBePISA server (https://www.ebi.ac.uk/pdbe/pisa/) and are colored in light green on chain A and light blue on chain C. Hydrogen bonds are established by residues F10, D11, K14 and Q19 on the α1a helix, and L55, D59 and I62 on the α3 helix. Hydrophobic interactions rely on residues I27, I30, I33, F34 and L37 on the α1b, and V102, L105, I106 and I109 on the α5 helix.

**Supplementary Figure S5.**
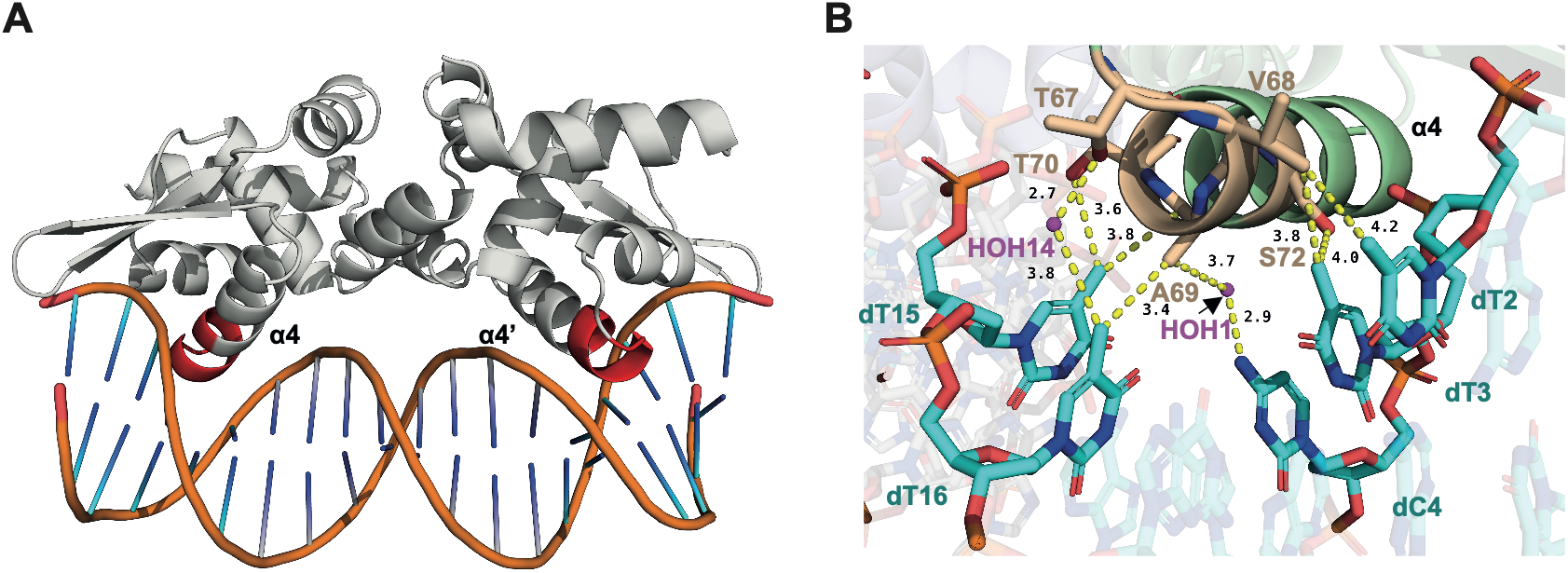
CadC-DNA complex. **A)** Cartoon representation of the CadC dimer bound to operator DNA. The protein engages DNA via a canonical winged helix–turn–helix (wHTH) architecture, with the recognition helix α4 highlighted in red. **B)** Close-up view of base-specific interactions involving residues V68, A69, T67, T70 and S72. Nucleotides from the 5′ and 3′ strands are coloured in cyan and labelled. Residues and water molecules involved in the interactions are shown in wheat and magenta, respectively. Dashed lines indicate the contact distances in angstroms.

**Supplementary Figure S6.**
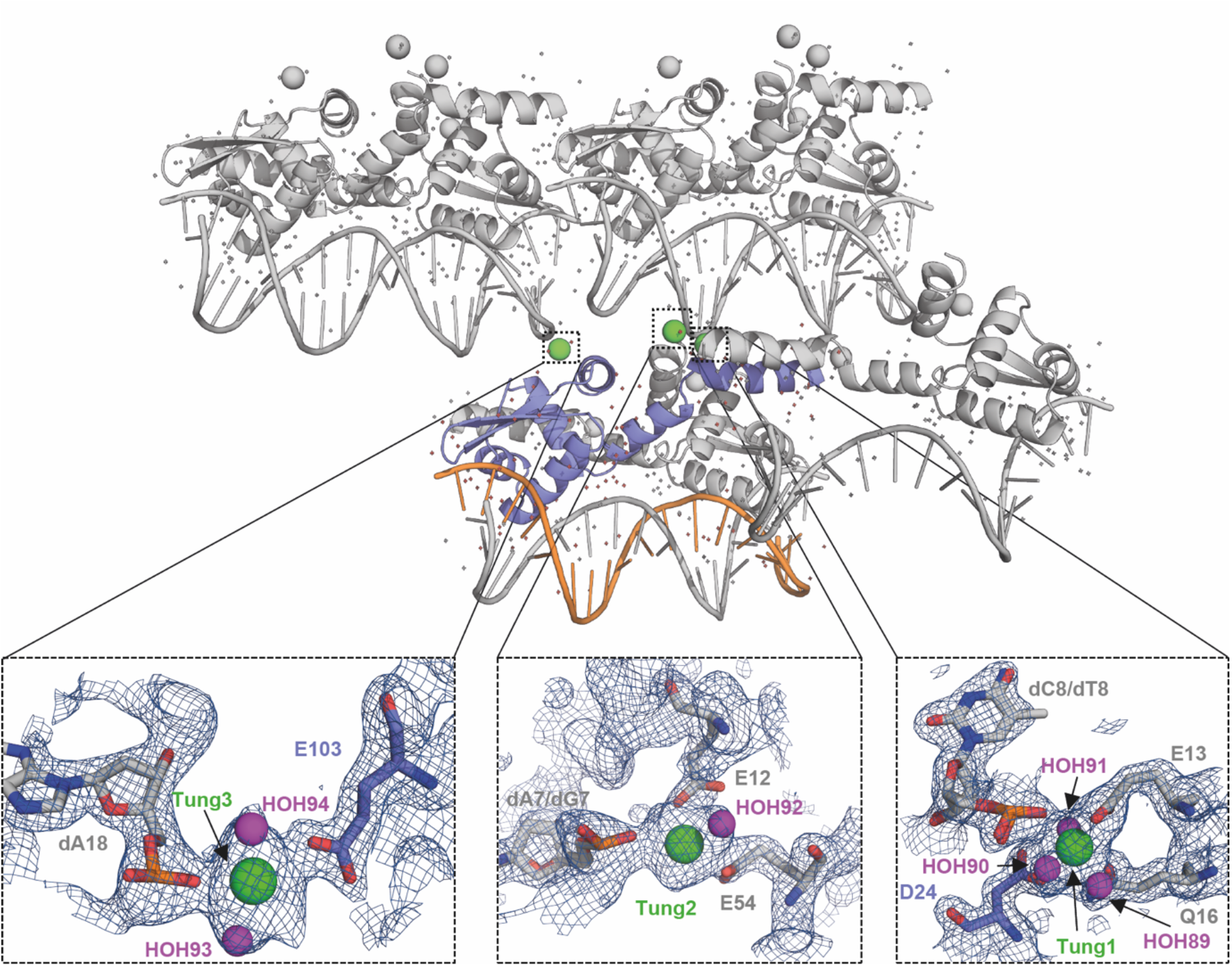
Structural overview of the CadC-DNA crystal packing and tungsten-binding sites. Top panel: Overall crystal packing arrangement showing the CadC-dsDNA complex in the asymmetric unit (AU), with the CadC protomer colored blue and its associated ssDNA colored orange. Symmetry-related CadC-DNA complexes are shown in grey. Tungsten atoms identified in the structure are represented as green spheres, whereas water molecules are shown as magenta spheres. The tungsten atoms are located at crystal-packing interfaces and are not associated with a conserved metal-binding pocket or a specific functional site, suggesting nonspecific binding arising from the crystallization conditions. Bottom panels: Close-up views of the three tungsten-binding sites (Tung1-Tung3). The 2Fo-Fc electron-density map is shown as a blue mesh contoured at 1.0σ around each site. Tungsten atoms are represented as green spheres and coordinating water molecules as magenta spheres. Residues, DNA nucleotides and solvent molecules involved in tungsten coordination are indicated.

**Supplementary Figure S7.**
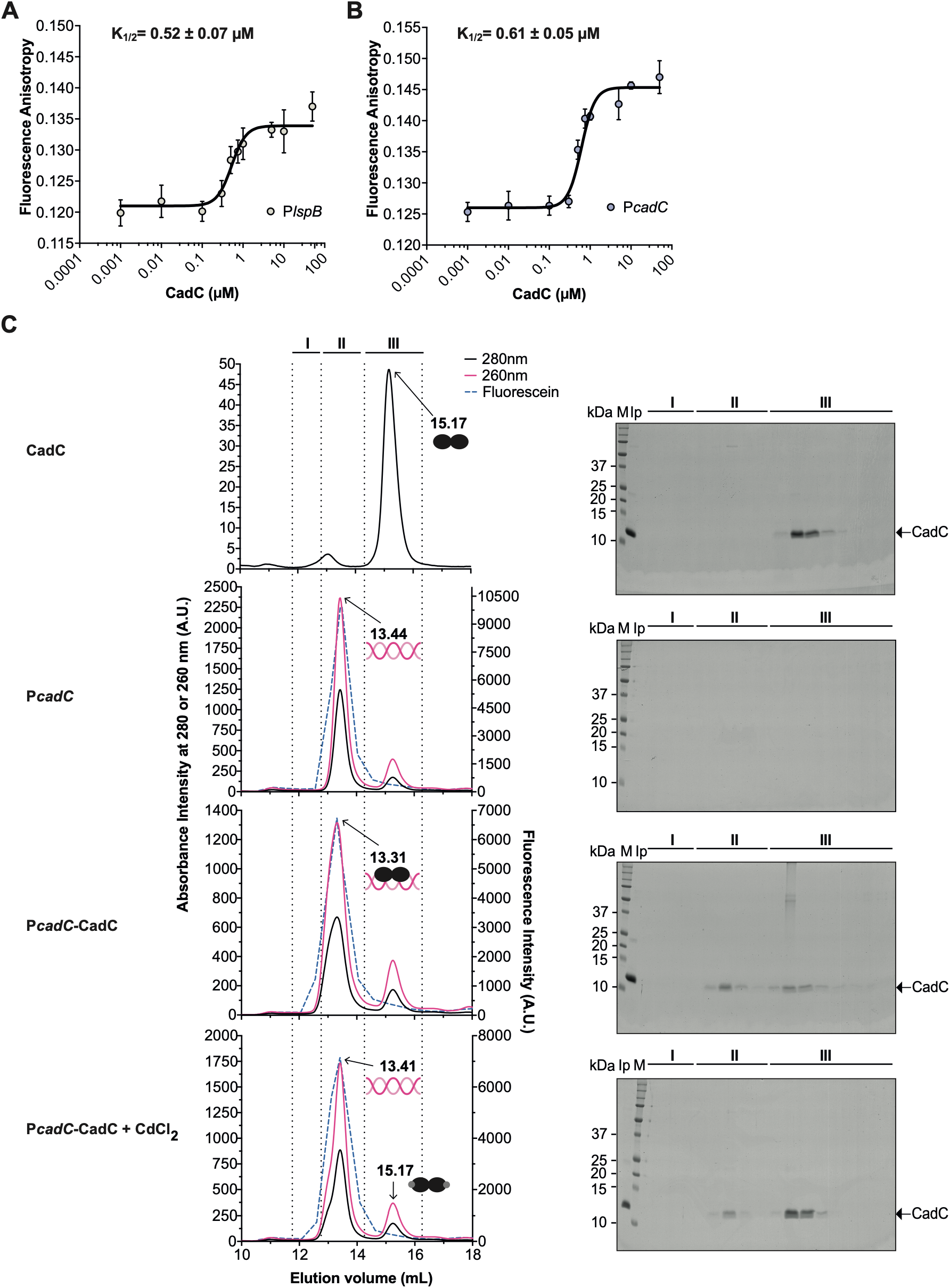
CadC binding to the CadC-Boxes in the *lspB* and *cadC* promoters. Fluorescence anisotropy analysis of CadC binding to the CadC-Box in the **(A)** *lspB* and **(B)** *cadC* promoters. Increasing concentrations of CadC (0–50 µM) were titrated at 22 °C into 5′-fluorescein (FAM)-labeled 33 bp oligonucleotides from P*lspB* or P*cadC*. Data represent the mean ± SD of three independent experiments. **(C)** Size exclusion chromatography profiles were monitored at 280 nm (CadC – black line) and 260 nm (P*cadC* – light red line). FAM fluorescence (light blue dashed line) was also measured for each collected fraction in a fluorescence plate reader. Chromatograms shown correspond to samples containing: CadC; FAM-P*cadC* alone, FAM-P*cadC* complexed with CadC in the absence of Cd^2+^, and CadC displaced from FAM-P*cadC* in the presence of Cd^2+^. Elution times for each peak are indicated above the chromatograms. Eluted fractions were labelled as I, II and III, and were analysed by SDS-PAGE electrophoresis, where the protein band corresponds to the monomeric CadC. Lane M, molecular mass markers (kDa; sizes shown on the left).

**Supplementary Figure S8.**
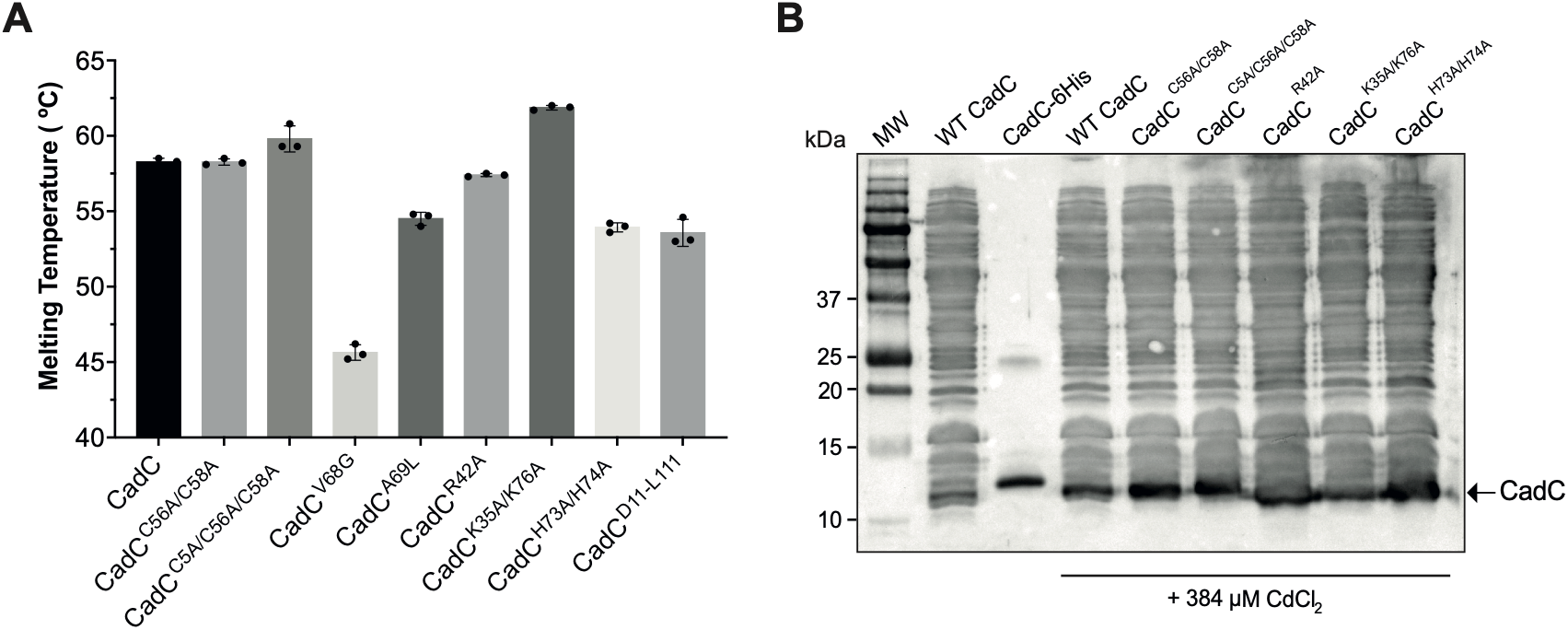
**(A)** Thermal shift assays performed with 5 µM of the indicated purified CadC variants. Protein unfolding was monitored with SYPRO Orange dye, and melting temperatures were determined from the inflection points of the unfolding curves. Data represent the mean ± SD of three independent experiments. **(B)** Western blot analysis of CadC expression in *Lm* mutant strains. Total cytosolic protein extracts from exponential-phase Δ*cadC* bacteria expressing the indicated CadC variants from a non-integrative plasmid grown in a chemically defined medium for 5 h in the absence or presence of 384 µM CdCl2, were analysed by immunoblot and CadC detected with an anti-CadC serum. Purified CadC (CadC-6His) was used as control.

**Supplementary Figure S9.**
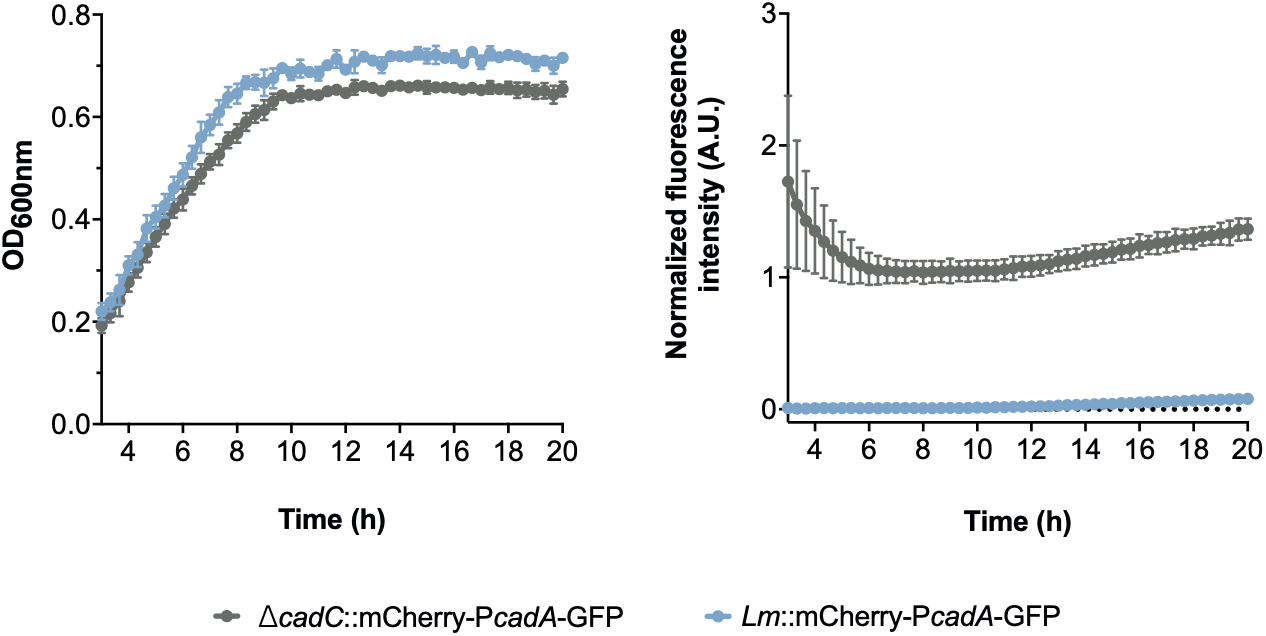
Growth of *Lm* strains constitutively expressing mCherry and carrying the P*cadA*-GFP (light blue line) transcriptional reporter fusions, as well as the Δ*cadC* mutant carrying the P*cadA*-GFP (gray line) transcriptional reporter fusion, were monitored in chemically defined medium. For all experiments, mid-log phase bacteria were diluted in a chemical defined medium (1:20) and bacterial growth was monitored by measuring the optical densities at 600 nm (OD600) every 20 min. Data represent the mean ± SD of three independent experiments.

**Supplementary Figure S10.**
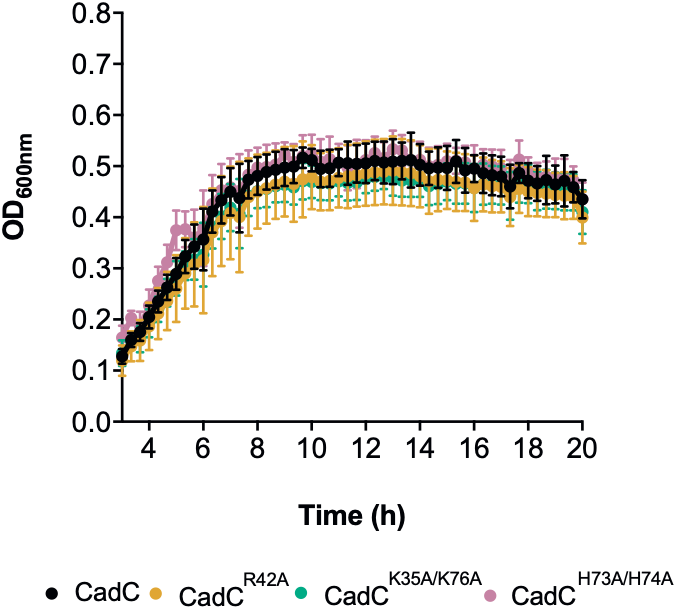
Mutations of CadC residues K35, R42, H73, H74 and K76 do not compromise bacterial growth. Bacterial growth (OD600) of *Lm* Δ*cadC* strains expressing CadC alanine substitutions (K35A, R42A, H73A/H74A, or K35A/K76A) and the GFP-P*cadA* transcriptional reporter fusion, was monitored in chemically defined medium. Mid-log-phase cultures were diluted (1:20) and grown at 37°C. Optical densities at 600 nm (OD600) were monitored every 20 min. Data represent the mean ± SD of three independent experiments.

**Supplementary Figure S11.**
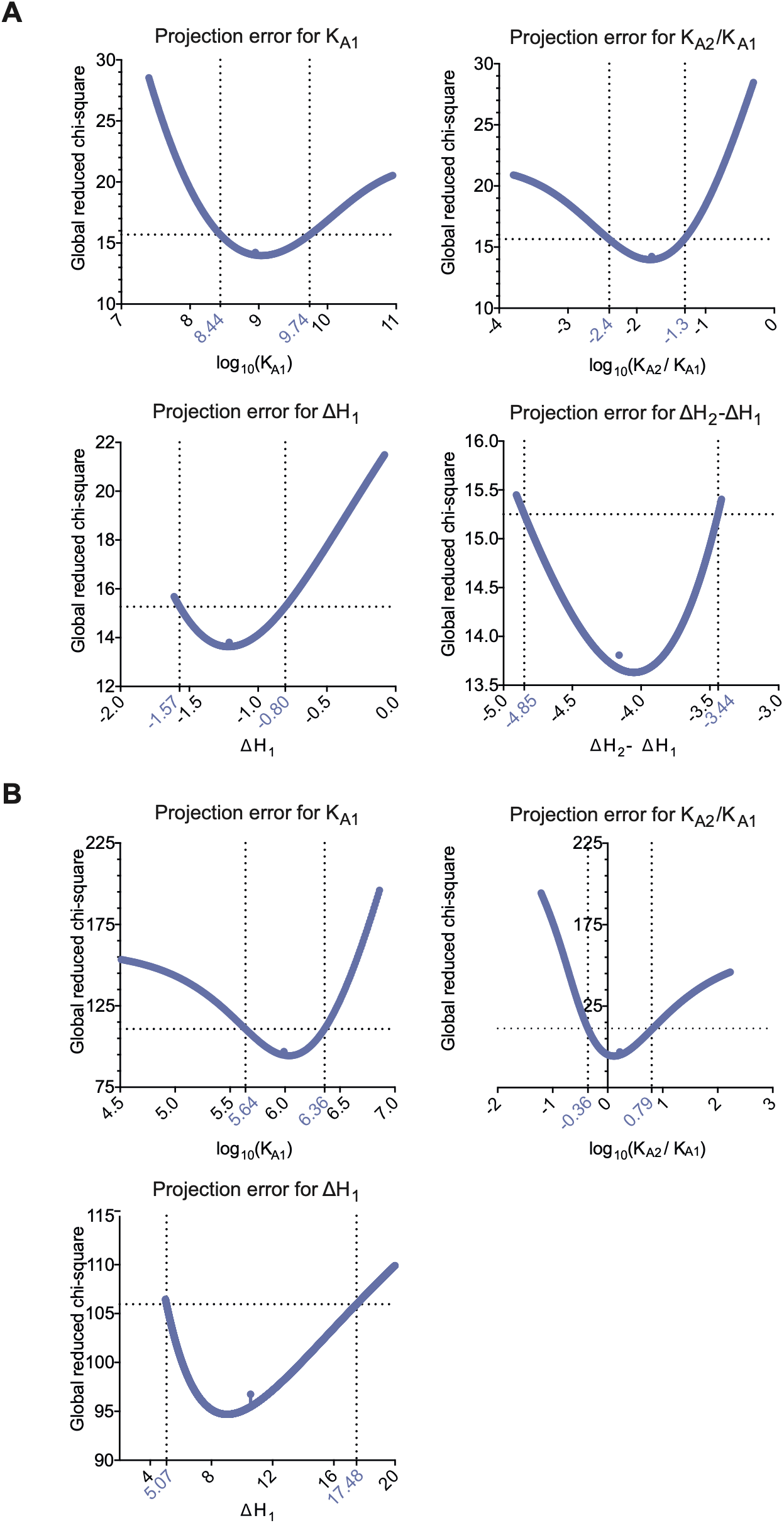
Two-dimensional error surface projections for the thermodynamic parameters determined by ITC. Projection error plots for (log10KA1), (log10KA2/KA1), (ΔH1) and (ΔH2-ΔH1) are shown. The minimum of each curve corresponds to the best-fit parameter value (minimum chi-square). Vertical dashed lines indicate the 68.3 % confidence interval limits and the horizontal dashed line represents the chi-square threshold corresponding to the 68.3 % confidence level. For the ZnCl2 titration, the projection error analysis did not yield a confidence interval for ΔH2-ΔH1.

**Supplementary Figure S12.**
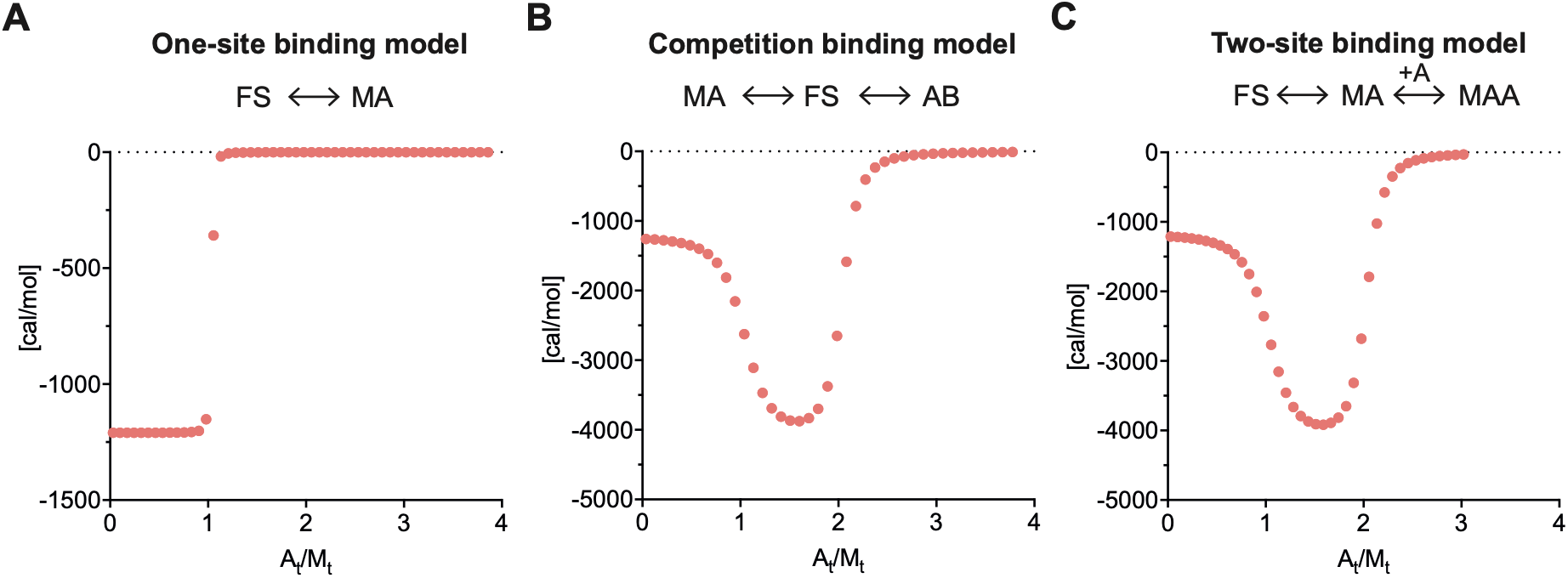
Simulated ITC binding isotherms for different binding models. Simulated calorimetric profiles were generated using AFFINImeter (1) for three binding models designed to illustrate how the presence of a high-affinity binding event together with a lower-affinity event can influence the shape of an ITC titration. For direct ITC measurements, very high C values (C= [Protein]/Kd) can result in sharply stoichiometric transitions that provide limited information for precise estimation of the corresponding binding constant. In the CadC-Cd²⁺ titrations, the protein concentration was 0.01 mM and the fitted high-affinity event had an apparent Kd of ∼ 1 nM, corresponding to C ∼ 10000. Under such conditions, competition approaches can broaden the binding transition and extend the experimentally informative range. The simulations below test whether the lower-affinity second binding event in CadC could produce an analogous broadening effect. **(A)** A simple one-site binding model (M+A⇌MA) in which the protein (M) is at 0.01 mM and contains a single binding site with Kd = 1 nM (Ka = 8.91×10^8^ M^-1^; ΔH = -1210 cal/mol). Ligand A was present in the syringe at 0.200 mM. At C ∼ 10 000, the simulated binding isotherm shows a sharp transition characteristic of near-stoichiometric binding, making the absolute affinity difficult to determine precisely from the direct titration alone. **(B)** A competition model (AB⇌B+M+A⇌MA) in which the cell additionally contains 0.01 mM of a competitor B with an affinity comparable to the lower-affinity CadC binding event (Kd=60 nM; Ka= 1.48×10^7^ M^-1^ and ΔH= -4160 cal/mol). The ligand concentration in the syringe was increased to 0.250 mM to account for binding to both protein and competitor. The competitor operates at C ∼ 167, producing a broader transition with measurable heat changes across the titration and illustrating how competition can increase the information content of the binding isotherm. **(C)** A two-site binding model (M+2A⇌MA+A⇌MA2) analogous to that used to describe Cd²⁺ binding to CadC. The first site has the same affinity and enthalpy as in **(A)** (Kd = 1nM), whereas the second site has the same affinity and enthalpy as the competitor in **(B)** (Kd = 60 nM). The resulting simulated isotherm is strongly broadened relative to the one-site model and resembles that obtained in the presence of an external competitor. This illustrates how a lower-affinity second binding event can broaden the calorimetric profile and thereby provide additional constraints when fitting a two-site model. These simulations support the plausibility of the fitted two-event model but do not independently validate the absolute value of the highest-affinity binding constant. In all simulations, the x-axis represents the molar ratio of metal (A) to protein (M) (At/Mt), and the y-axis shows the integrated heat per injection (cal/mol). Simulated titrations consisted of 40 injections comprising one initial 2 μL injection followed by 39 injections of 5 μL. FS denotes the free species.

**Supplementary Figure S13.**
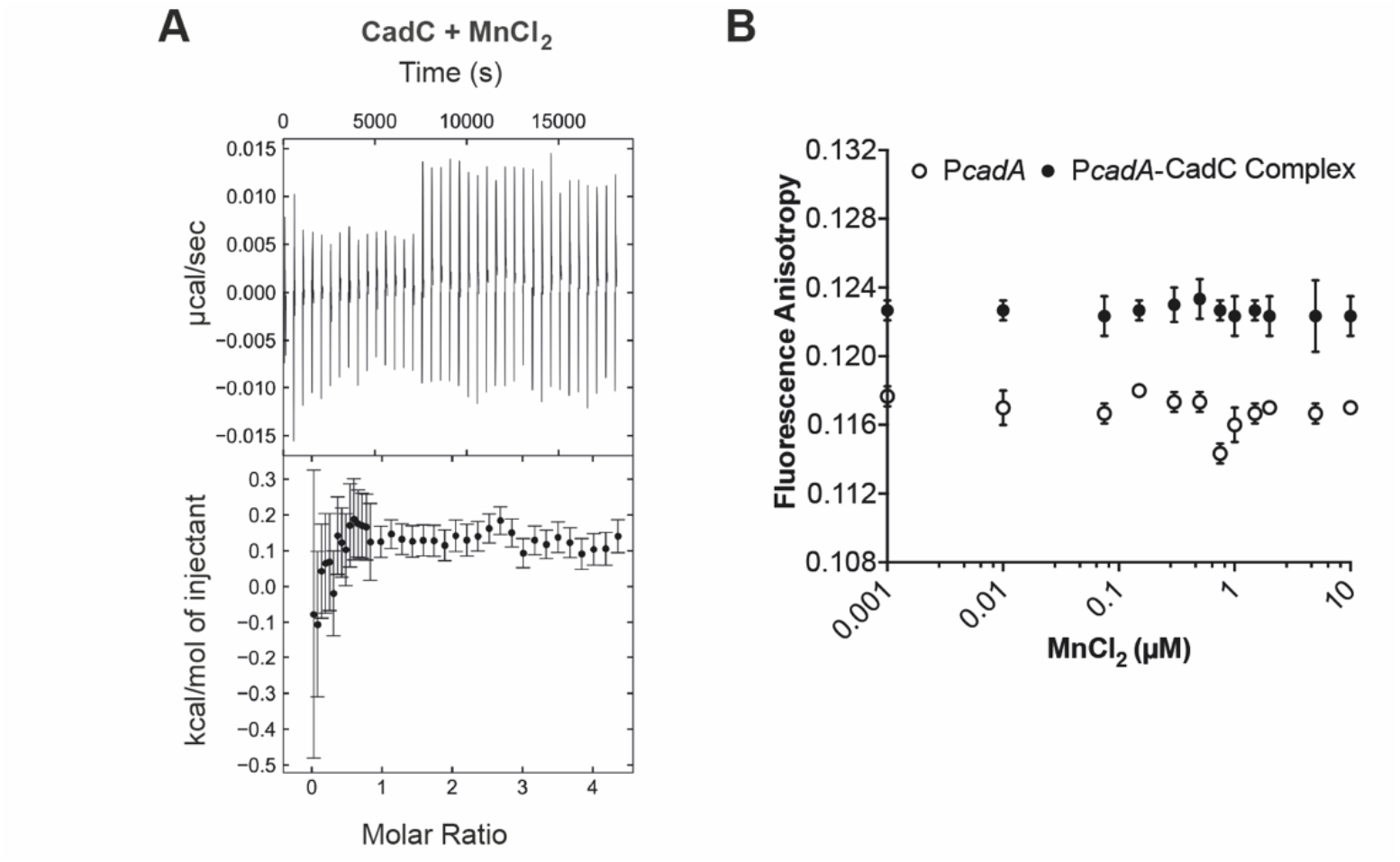
**(A)** ITC measurements of the interaction of CadC (10 µM) with MnCl2 (200 µM) at 15°C. Upper panels show raw data plotted as power over time, and lower panels show the integrated, concentration normalized ITC data. Data shown are representative of two independent experiments yielding similar results. **(B)** Fluorescence anisotropy analysis of the apo-CadC-P*cadA* complex titrated with increasing concentrations of MnCl2 (0-10 µM) at 22°C (black dots). P*cadA* alone (empty circle dots) was included as a negative control. Data represent the mean ± SD of three independent experiments.

**Supplementary Figure S14.**
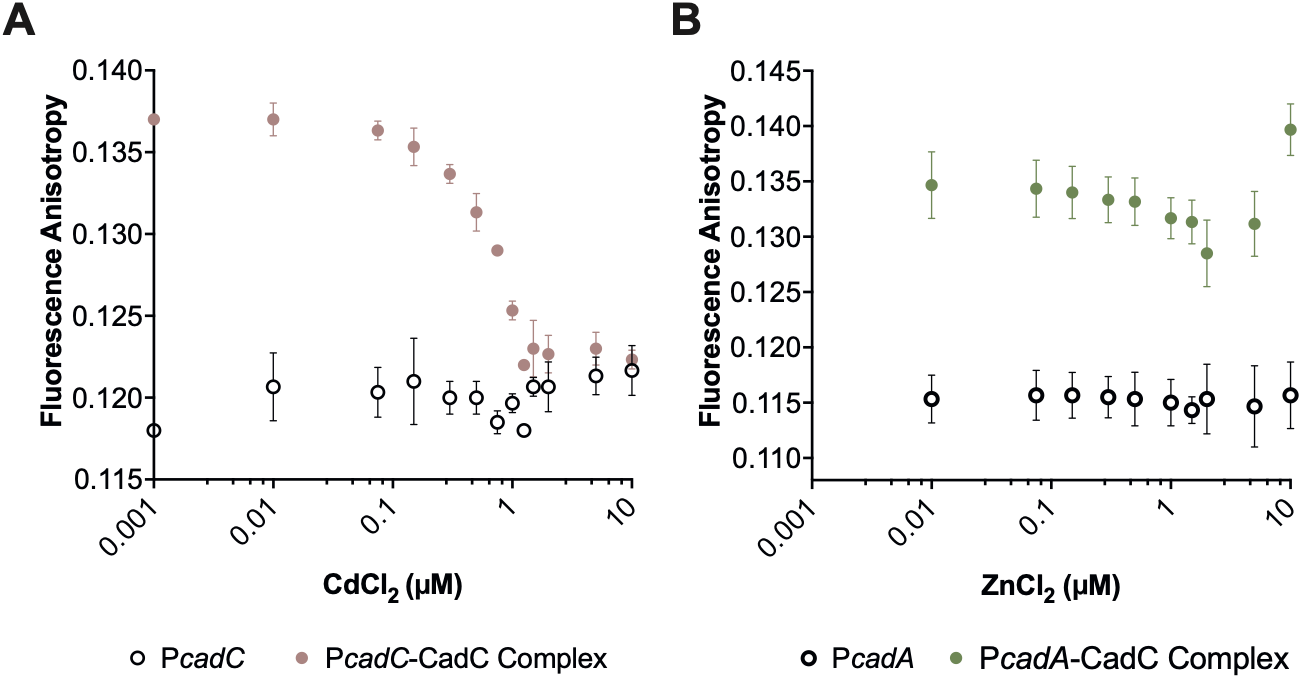
Cd^2+^, but not Zn^2+^, disrupts CadC-DNA complexes. Fluorescence anisotropy analysis of CadC-DNA complexes in response to metal ion titration. **(A)** CadC-P*cadC* complex titrated with increasing concentrations CdCl2 (0-10 µM) at 22 °C (red circles). **(B)** CadC-P*cadA* complex titrated with increasing concentrations of ZnCl2 (green circles). FAM-P*cadA* alone (open circles) was included as negative control. Data represent the mean ± SD of three independent experiments.

**Supplementary Figure S15.**
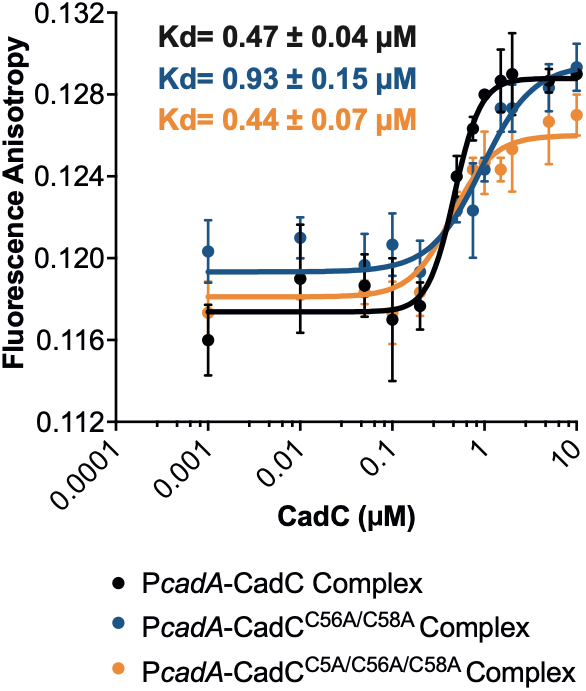
CadC mutated in coordinator cysteines retains its ability to complex with a DNA sequence containing a CadC box (e.g. P*cadA)*. Fluorescence anisotropy of FAM-P*cadA* titrated with increasing concentrations of CadC (black line), CadC^C56A/C58A^ (red line) or CadC^C5A/C56A/C58A^ (blue line) (0-10 µM) at 22°C. Data represent the mean ± SD of three independent experiments.

**Supplementary Figure S16.**
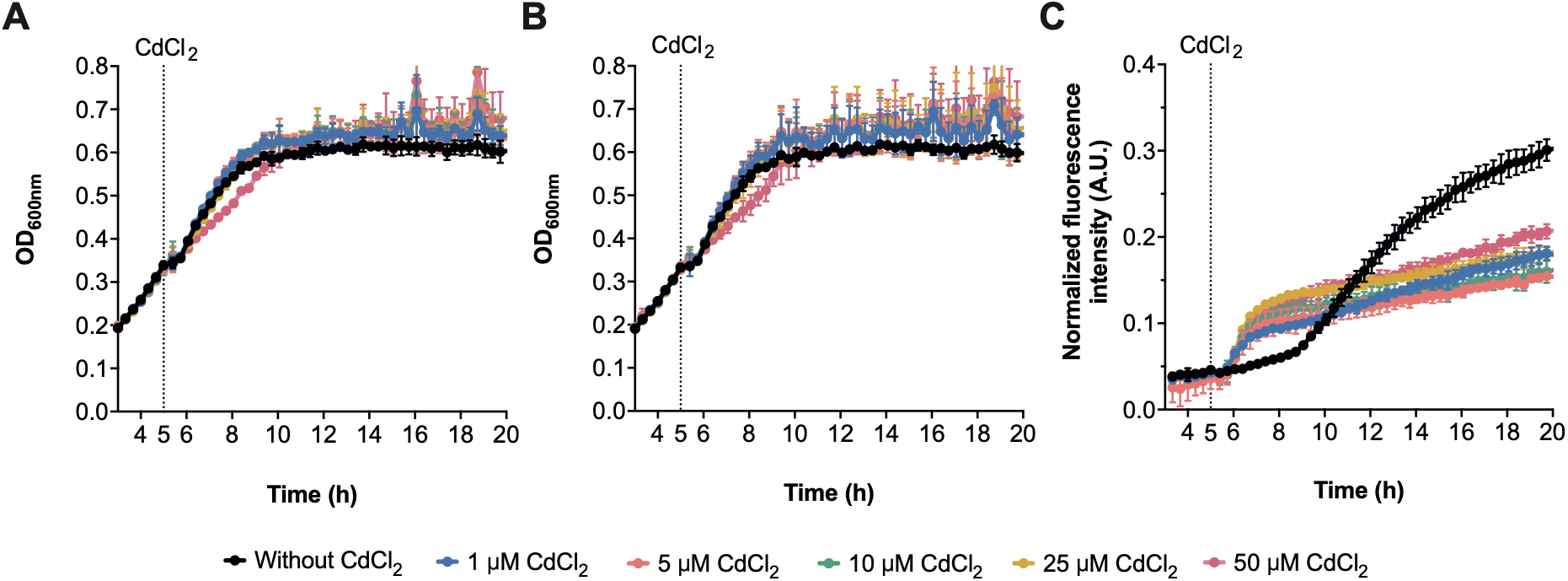
Cadmium induces *cadC* derepression without affecting bacterial growth. The growth of wild-type *Lm* constitutively expressing mCherry and carrying the **(A)** P*cadA*-GFP or **(B)** P*cadC-*GFP transcriptional reporter fusions was monitored in chemically defined medium. **(C)** Cadmium induces derepression of *cadC in vivo*. GFP fluorescence of wild-type *Lm* carrying the GFP-*cadC* promoter transcriptional reporter fusion, supplemented or not with CdCl2. For all experiments, mid-log phase bacteria were diluted in a chemically defined medium (1:20) and increasing concentrations of CdCl2 (1, 5, 10, 25 and 50 µM) were added after 5 h of growth (dashed line). Optical densities at 600 nm (OD600) were monitored every 20 min. Data represent the mean ± SD of three independent experiments.

**Supplementary Figure S17.**
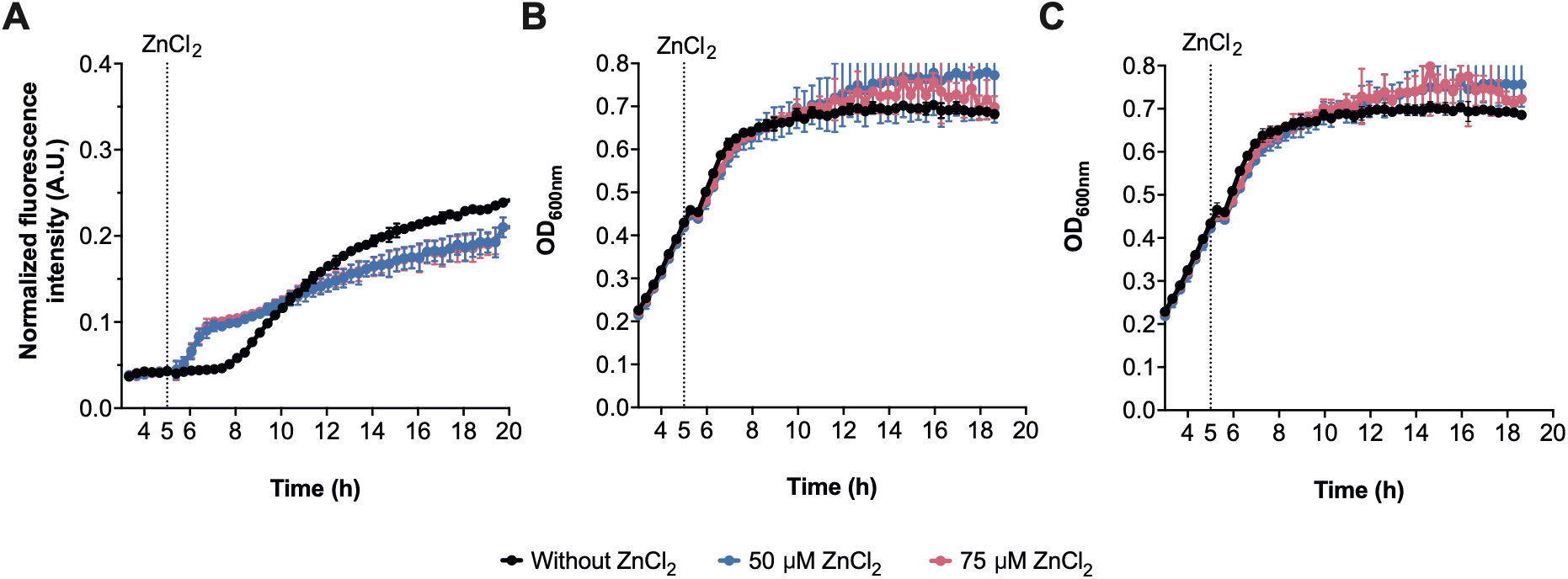
**(A)** Zinc induces derepression of *cadC in vivo*. GFP fluorescence of *Lm* carrying the P*cadC*-GFP transcriptional reporter fusion, grown in chemically defined medium in the absence of ZnCl2 (black) or supplemented with 50 µM (blue) or 75 µM (red) ZnCl2 after 5 h of growth (dashed line). **(B-C)** The growth of wild-type *Lm* constitutively expressing mCherry and carrying the **(B)** P*cadC*-GFP or **(C)** P*cadA*-GFP transcriptional reporter fusions were monitored in chemically defined medium. For all experiments, mid-log phase bacteria were diluted in a chemically defined medium (1:20) and increasing concentrations of ZnCl2 (50 and 75 µM) were added after 5 h of growth (dashed line). Optical densities at 600 nm (OD600) were monitored every 20 min. Data represent the mean ± SD of three independent experiments.

**Supplementary Figure S18.**
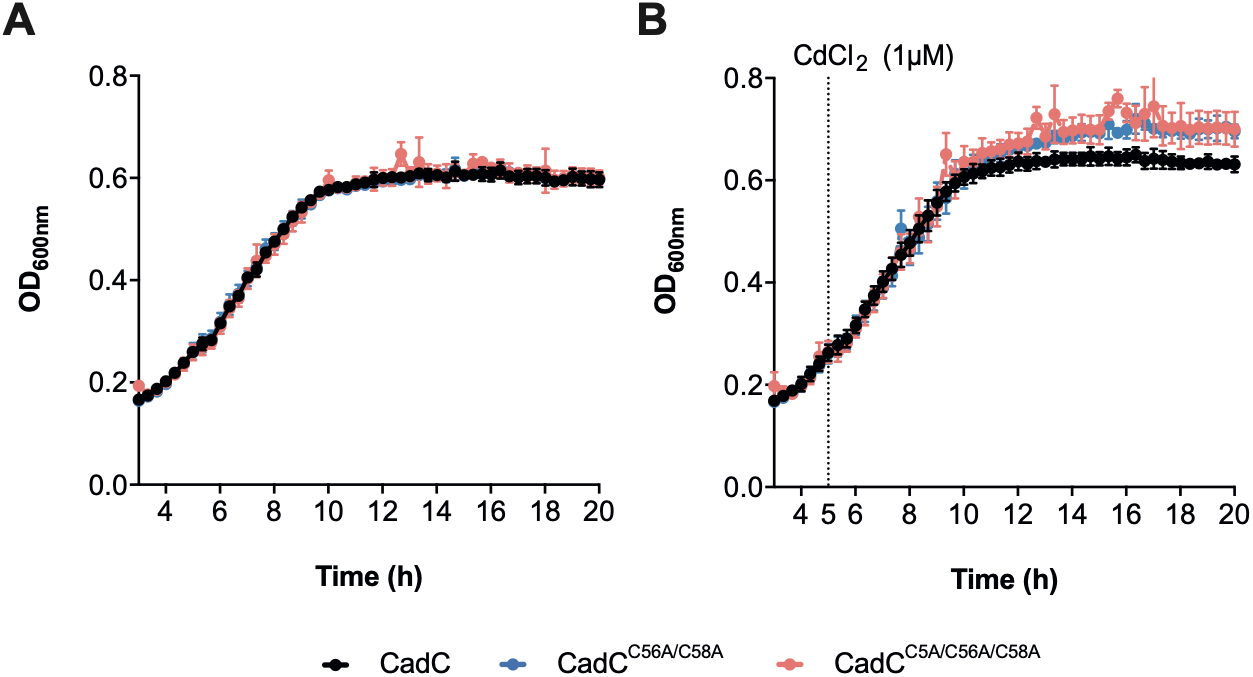
Cysteine mutations do not compromise bacterial growth. Bacterial growth (OD600) of *Lm* Δ*cadC* carrying the P*cadA*-GFP transcriptional reporter fusion and expressing wild-type CadC or CadC cysteine mutants (CadC^C56A/C58A^ and CadC^C5A/C56A/C58A^) was monitored in chemically defined medium. Mid-log-phase cultures were diluted (1:20) and grown either **(A)** in the absence of cadmium or **(B)** supplemented with 1 µM CdCl2 after 5 h of growth (dashed line).

**Supplementary Table 1.**
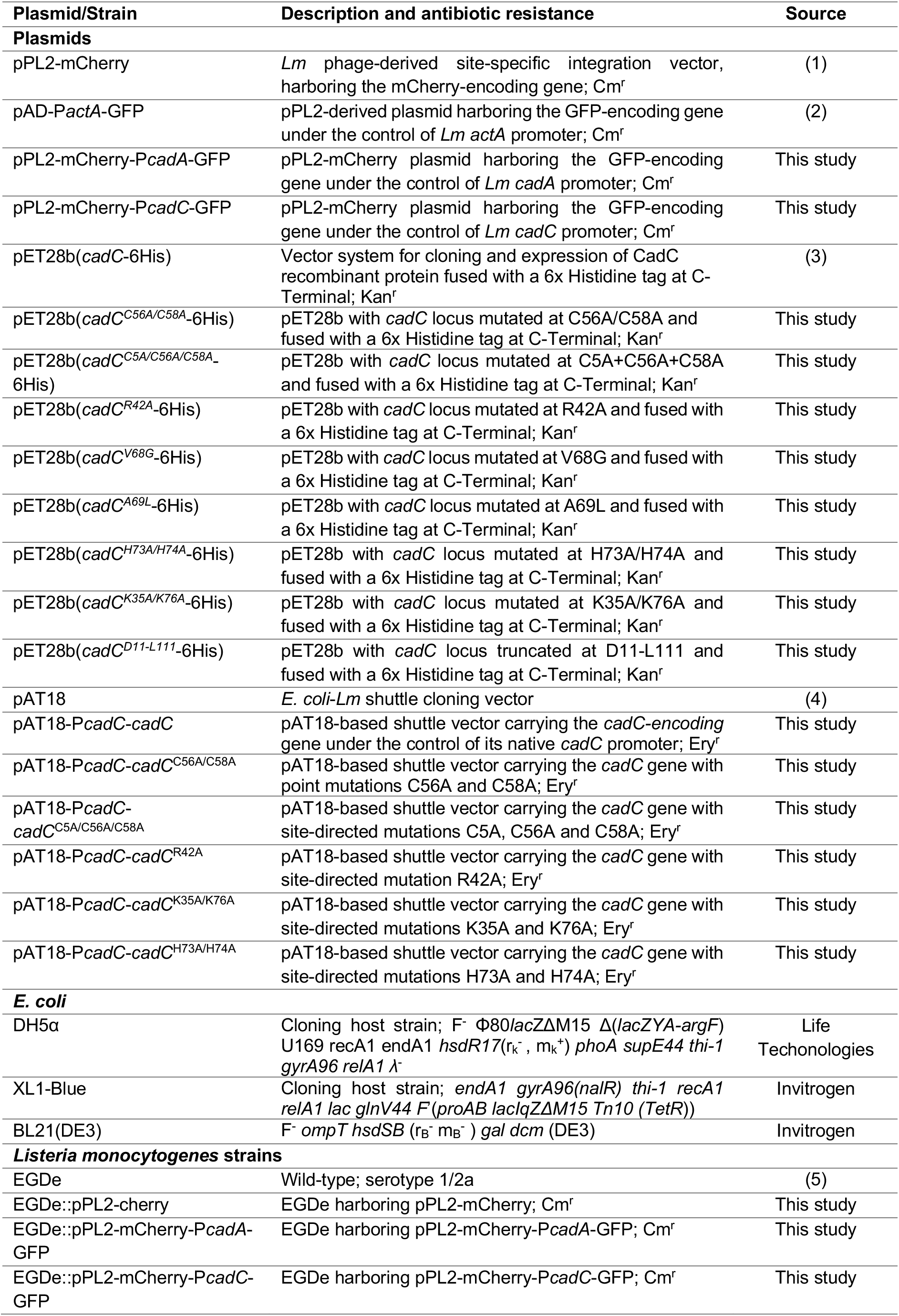
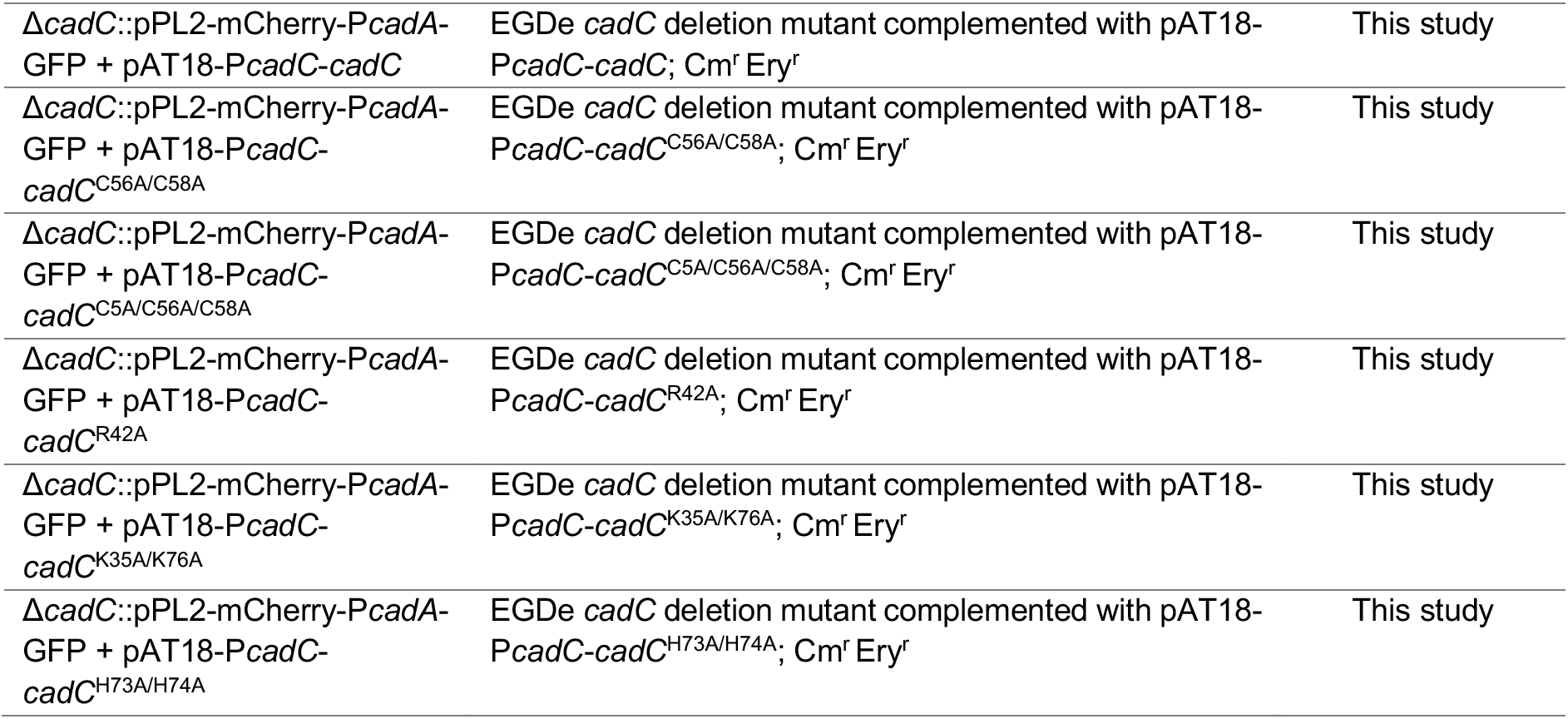
Plasmids and bacterial strains.

**Supplementary Table 2.**
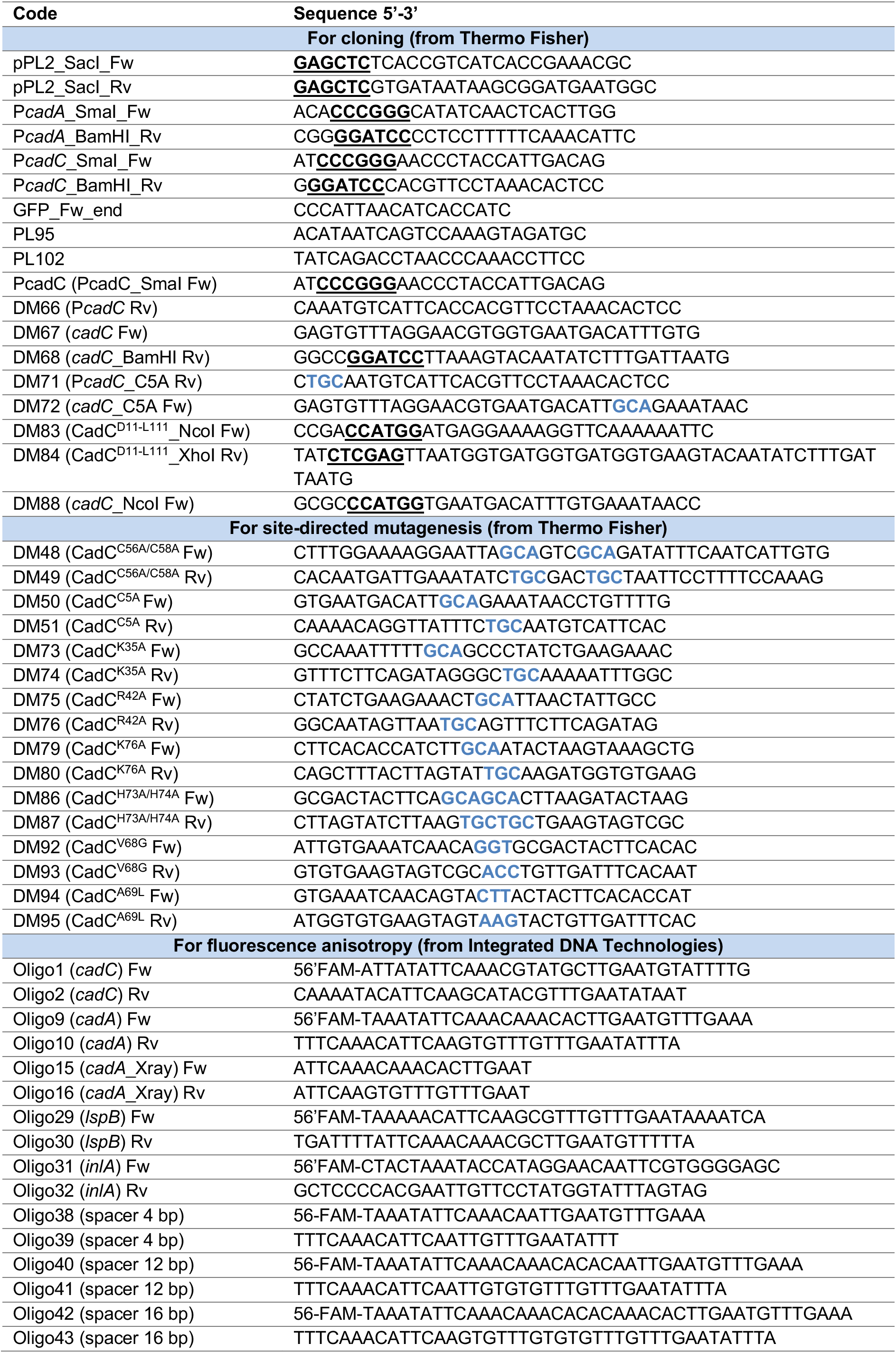
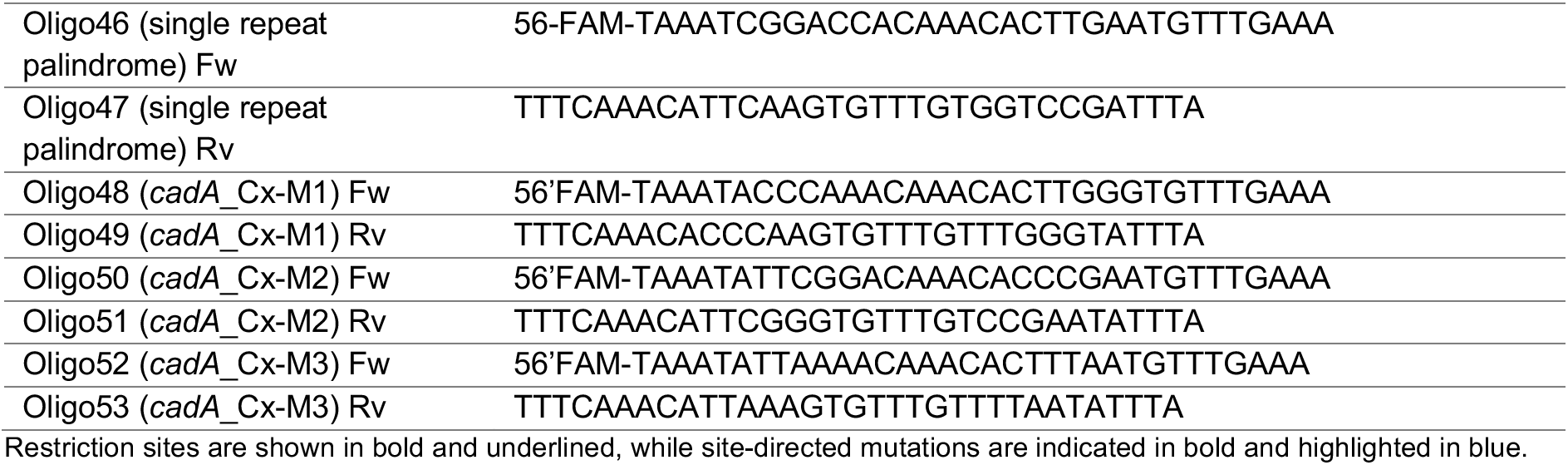
Oligonucleotides used in this study.

**Supplementary Table 3.**
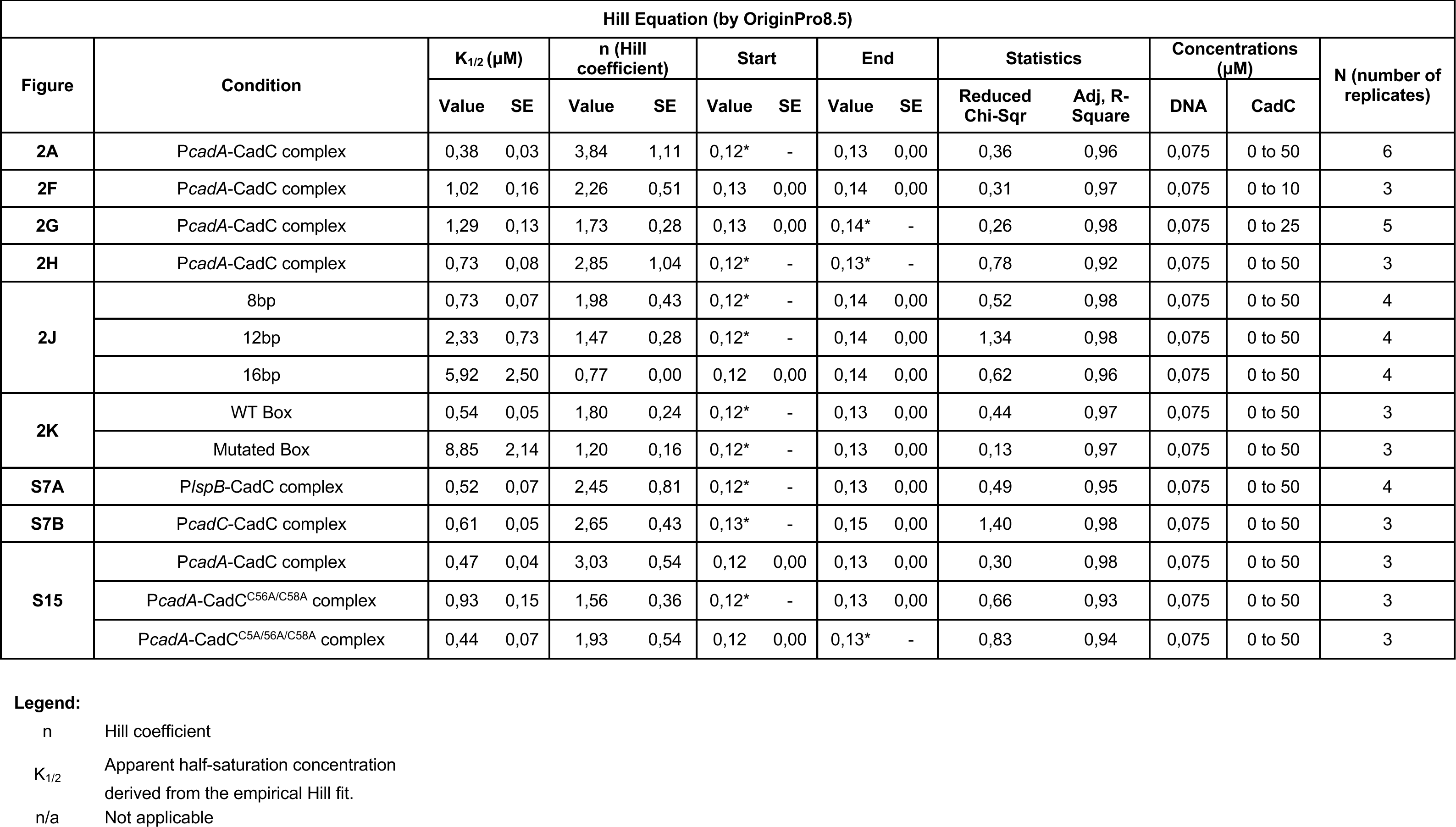
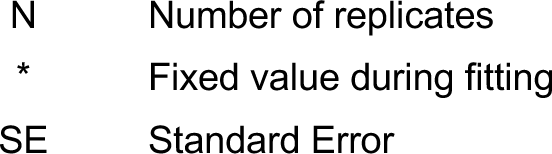
Summary of Hill-model apparent parameters describing Protein-DNA binding interactions.

| Hill Equation (by OriginPro8.5) |  |  |  |  |  |  |  |  |  |  |  |  |  |  |
| --- | --- | --- | --- | --- | --- | --- | --- | --- | --- | --- | --- | --- | --- | --- |
| Figure | Condition | K<sub>1/2</sub> (μM) |  | n (Hill coefficient) |  | Start |  | End |  | Statistics |  | Concentrations (μM) |  | N (number of replicates) |
|  |  | Value | SE | Value | SE | Value | SE | Value | SE | Reduced Chi-Sqr | Adj, R-Square | DNA | CadC |  |
| 2A | PcadA-CadC complex | 0,38 | 0,03 | 3,84 | 1,11 | 0,12* | - | 0,13 | 0,00 | 0,36 | 0,96 | 0,075 | 0 to 50 | 6 |
| 2F | PcadA-CadC complex | 1,02 | 0,16 | 2,26 | 0,51 | 0,13 | 0,00 | 0,14 | 0,00 | 0,31 | 0,97 | 0,075 | 0 to 10 | 3 |
| 2G | PcadA-CadC complex | 1,29 | 0,13 | 1,73 | 0,28 | 0,13 | 0,00 | 0,14* | - | 0,26 | 0,98 | 0,075 | 0 to 25 | 5 |
| 2H | PcadA-CadC complex | 0,73 | 0,08 | 2,85 | 1,04 | 0,12* | - | 0,13* | - | 0,78 | 0,92 | 0,075 | 0 to 50 | 3 |
| 2J | 8bp | 0,73 | 0,07 | 1,98 | 0,43 | 0,12* | - | 0,14 | 0,00 | 0,52 | 0,98 | 0,075 | 0 to 50 | 4 |
|  | 12bp | 2,33 | 0,73 | 1,47 | 0,28 | 0,12* | - | 0,14 | 0,00 | 1,34 | 0,98 | 0,075 | 0 to 50 | 4 |
|  | 16bp | 5,92 | 2,50 | 0,77 | 0,00 | 0,12 | 0,00 | 0,14 | 0,00 | 0,62 | 0,96 | 0,075 | 0 to 50 | 4 |
| 2K | WT Box | 0,54 | 0,05 | 1,80 | 0,24 | 0,12* | - | 0,13 | 0,00 | 0,44 | 0,97 | 0,075 | 0 to 50 | 3 |
|  | Mutated Box | 8,85 | 2,14 | 1,20 | 0,16 | 0,12* | - | 0,13 | 0,00 | 0,13 | 0,97 | 0,075 | 0 to 50 | 3 |
| S7A | PlspB-CadC complex | 0,52 | 0,07 | 2,45 | 0,81 | 0,12* | - | 0,13 | 0,00 | 0,49 | 0,95 | 0,075 | 0 to 50 | 4 |
| S7B | PcadC-CadC complex | 0,61 | 0,05 | 2,65 | 0,43 | 0,13* | - | 0,15 | 0,00 | 1,40 | 0,98 | 0,075 | 0 to 50 | 3 |
| S15 | PcadA-CadC complex | 0,47 | 0,04 | 3,03 | 0,54 | 0,12 | 0,00 | 0,13 | 0,00 | 0,30 | 0,98 | 0,075 | 0 to 50 | 3 |
|  | PcadA-CadC<sup>C56A/C58A</sup> complex | 0,93 | 0,15 | 1,56 | 0,36 | 0,12* | - | 0,13 | 0,00 | 0,66 | 0,93 | 0,075 | 0 to 50 | 3 |
|  | PcadA-CadC<sup>C5A/56A/C58A</sup> complex | 0,44 | 0,07 | 1,93 | 0,54 | 0,12 | 0,00 | 0,13* | - | 0,83 | 0,94 | 0,075 | 0 to 50 | 3 |

- n Hill coefficient - K<sub>1/2</sub> Apparent half-saturation concentration derived from the empirical Hill fit. - n/a Not applicable
N Number of replicates \* Fixed value during fitting SE Standard Error

**Supplementary Table 4.**
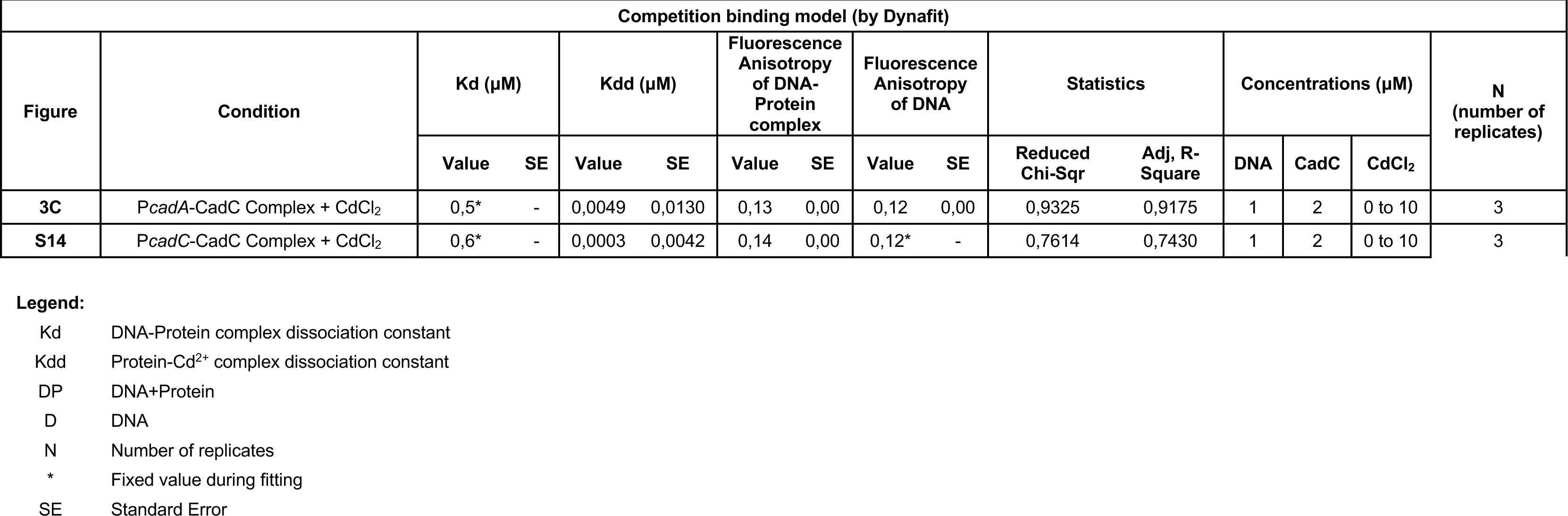
Summary of DynaFit-derived apparent parameters from competition binding analysis.

| Competition binding model (by Dynafit) |  |  |  |  |  |  |  |  |  |  |  |  |  |  |  |
| --- | --- | --- | --- | --- | --- | --- | --- | --- | --- | --- | --- | --- | --- | --- | --- |
| Figure | Condition | Kd (μM) |  | Kdd (μM) |  | Fluorescence Anisotropy of DNA-Protein complex |  | Fluorescence Anisotropy of DNA |  | Statistics |  | Concentrations (μM) |  |  | N (number of replicates) |
|  |  | Value | SE | Value | SE | Value | SE | Value | SE | Reduced Chi-Sqr | Adj, R-Square | DNA | CadC | CdCl<sub>2</sub> |  |
| 3C | PcadA-CadC Complex + CdCl<sub>2</sub> | 0,5* | - | 0,0049 | 0,0130 | 0,13 | 0,00 | 0,12 | 0,00 | 0,9325 | 0,9175 | 1 | 2 | 0 to 10 | 3 |
| S14 | PcadC-CadC Complex + CdCl<sub>2</sub> | 0,6* | - | 0,0003 | 0,0042 | 0,14 | 0,00 | 0,12* | - | 0,7614 | 0,7430 | 1 | 2 | 0 to 10 | 3 |

Kd DNA-Protein complex dissociation constant Kdd Protein-Cd<sup>2+</sup> complex dissociation constant DP DNA+Protein D DNA N Number of replicates \* Fixed value during fitting SE Standard Error

